# Human, Nonhuman Primate, and Bat Cells Are Broadly Susceptible to Tibrovirus Particle Cell Entry

**DOI:** 10.1101/507350

**Authors:** Yíngyún Caì, Shuĭqìng Yú, Rohit K. Jangra, Elena N. Postnikova, Jiro Wada, Robert B. Tesh, Sean P. J. Whelan, Michael Lauck, Michael R. Wiley, Courtney L. Finch, Sheli R. Radoshitzky, David H. O’Connor, Gustavo Palacios, Kartik Chandran, Charles Y. Chiu, Jens H. Kuhn

**Affiliations:** Integrated Research Facility at Fort Detrick, National Institute of Allergy and Infectious Diseases, National Institutes of Health, Fort Detrick, Frederick, MD, USA; Department of Microbiology and Immunology, Albert Einstein College of Medicine, Bronx, NY, USA; Department of Pathology, Center for Biodefense and Emerging Infectious Diseases, University of Texas Medical Branch, Galveston, TX, USA; Department of Microbiology and Immunobiology, Harvard Medical School, Boston, MA, USA; Department of Pathology and Laboratory Medicine, University of Wisconsin, Madison WI, USA; United States Army Medical Research Institute of Infectious Diseases, Fort Detrick, Frederick, MD, USA; University of California, San Francisco, CA, USA

**Keywords:** Bas-Congo virus, *Mononegavirales*, mononegavirus, *Rhabdoviridae*, rhabdovirus, tibrovirus, tropism, viral hemorrhagic fever

## Abstract

In 2012, the genome of a novel rhabdovirus, Bas-Congo virus, was discovered in the acute-phase serum of a Congolese patient with presumed viral hemorrhagic fever. In the absence of a replicating virus isolate, fulfilling Koch’s postulates to determine whether Bas-Congo virus is indeed a human virus and/or pathogen has been impossible. However, experiments with vesiculoviral particles pseudotyped with Bas-Congo glycoprotein suggested that Bas-Congo virus particles can enter cells from multiple animals, including humans. In 2015, genomes of two related viruses, Ekpoma virus 1 and Ekpoma virus 2, were detected in human sera in Nigeria. Isolates could not be obtained. Phylogenetic analyses led to the classification of Bas-Congo virus, Ekpoma virus 1, and Ekpoma virus 2 in the same genus, *Tibrovirus*, together with five biting midge-borne rhabdoviruses (i.e., Beatrice Hill virus, Bivens Arm virus, Coastal Plains virus, Sweetwater Branch virus, and Tibrogargan virus) not known to infect humans. Using individual recombinant vesiculoviruses expressing the glycoproteins of all eight known tibroviruses and more than 75 cell lines representing different animal species, we demonstrate that the glycoproteins of all tibroviruses can mediate vesiculovirus particle entry into human, bat, nonhuman primate, cotton rat, boa constrictor, and Asian tiger mosquito cells. Using four of five isolated authentic tibroviruses (i.e., Bivens Arm virus, Coastal Plains virus, Sweetwater Branch virus, and Tibrogargan virus), our experiments indicate that many cell types may be partially resistant to tibrovirus replication after virion cell entry. Consequently, experimental data solely obtained from experiments using tibrovirus surrogate systems (e.g., vesiculoviral pseudotypes, recombinant vesiculoviruses) cannot be used to predict whether Bas-Congo virus, or any other tibrovirus, infects humans.

## INTRODUCTION

The viral order *Mononegavirales* currently includes eleven families for negative-sense single-stranded RNA viruses (Maes et al., 2019). With 18 included genera, the family *Rhabdoviridae* is the largest and most diverse of the mononegaviral families (Walker et al., 2018; Maes et al., 2019). Yet, viruses of most genera are undercharacterized, and their potential as human pathogens remains largely unknown. This undercharacterization holds true, for instance, for the rhabdovirus genus *Tibrovirus* (Bourhy et al., 2005; Gubala et al., 2011), which was suspected to harbor only viruses without any clinical or veterinary significance. However, the description of a tibrovirus associated with suspected viral hemorrhagic fever in humans in 2012 challenged this assumption (Grard et al., 2012; Chiu et al., 2013).

The prototypical tibroviruses are Tibrogargan virus (TIBV, species *Tibrogargan tibrovirus*), Coastal Plains virus (CPV, species *Coastal Plains tibrovirus*), and Bivens Arm virus (BAV, species *Tibrogargan tibrovirus*) (Walker et al., 2018). TIBV was first described in 1980 as a rhabdovirus infecting biting midges (*Culicoides brevitaris*) that were collected around Peachester, Queensland, Australia, close to Mount Tibrogargan (Cybinski et al., 1980). Anti-TIBV antibodies were found in healthy cattle throughout Australia, in New Guinea, and in the US. Antibodies were also found in healthy Australian water buffalo and a Floridian white-tailed deer, but not in Australian camels, humans, dogs, goats, horses, pigs, sheep, wallabies, and possums, (Cybinski et al., 1980; Gibbs et al., 1989; Calisher et al., 1993; Gubala et al., 2011). Finally, subclinical TIBV infection was demonstrated in healthy sentinel cattle in Peachester (St George, 1985).

CPV was isolated in 1981 at Coastal Plains Research Station (today Coastal Plains Research Farm), Northern Territory, Australia, from a viremic but healthy steer. Anti-CPV antibodies were detected in healthy Australian buffalo, cattle, dogs, and a horse, but not in deer, humans, pigs, or wallabies. Anti-CPV antibodies were also found in healthy cattle from Papua New Guinea (Cybinski and Gard, 1986; Gard et al., 1988).

BAV was isolated in 1982 from biting midges (*Culicoides insignis*) collected near water buffaloes in Florida, USA, and anti-BAV antibodies were detected in healthy Floridian cattle, one horse and white-tailed deer, but not in sheep or wildebeest (Gibbs et al., 1989). Anti-BAV antibodies were also detected in healthy Trinidadian water buffaloes (Calisher et al., 1993) and cattle from Puerto Rico and St. Croix, United States Virgin Islands (Tuekam et al., 1991).

BAV, CPV, and TIBV produce viral particles with rhabdovirion-characteristic bullet-like morphologies (Cybinski et al., 1980; Cybinski and Gard, 1986; Gibbs et al., 1989). However, analysis of the complete genome sequences of CPV (13,203 nt) and TIBV (13,298 nt) revealed a unique rhabdovirus genome organization (*3’N-P-M-U1-U2-G-U3-L5’*) characterized by two novel genes of unknown function (*U1, U2*) located between the matrix protein (M) gene and the glycoprotein (G) gene and one gene of unknown function (U3) between the *G* gene and RNA-dependent RNA polymerase (L) gene. Each of these genes is defined as an independent transcriptional unit bounded by consensus transcription initiation and transcription termination/polyadenylation sequences. The TIBV genome differs from the CPV genome by the presence of a fourth unique open reading frame (U4 or Gx) overlapping the *G* gene (Gubala et al., 2011; Walker et al., 2015).

In recent years, the genus *Tibrovirus* has grown steadily. Most notably, Bas-Congo virus (BASV) was identified as a tibrovirus (Walker et al., 2015). BASV was detected by next generation sequencing (NGS) in an acute-phase serum sample from a human with suspected viral hemorrhagic fever in Mangala, Bas-Congo Province (today Kongo Central Province), Democratic Republic of the Congo (Grard et al., 2012). Unfortunately, a BASV isolate could not be obtained. Therefore, whether BASV indeed infects humans or causes disease remains unclear. A recent *in silico* analysis of the BASV genome using a novel machine learning algorithm indicates that the natural host of BASV is an artiodactyl and that BASV may be vectored by biting midges (Babayan et al., 2018). The BASV genomic sequence (11,892 nt) remains incomplete: the sequences of all genes have been obtained except those of the *N* and *L* genes, which are incomplete at their extreme termini (Grard et al., 2012). Hence, a reverse genetics system to rescue replicating BASV could not yet be established and the question of BASV host tropism can therefore only be examined using indirect means.

Genomes of another two tibroviruses, Ekpoma virus 1 (EKV-1, 12,659 nt) and Ekpoma virus 2 (EKV-2, 12,674 nt), were discovered by NGS in blood samples from apparently healthy humans in Nigeria (Stremlau et al., 2015). In addition, an EKV-2 like genome detected in a human from Angola was recently deposited in GenBank (accession #MF079256; 12,638 nt) but remains to be described. As in the case of BASV, cell-culture isolates for these viruses are not available, their genome sequences are incomplete at their termini (Stremlau et al., 2015), and whether any of these virus actually infects humans, or is the cause of any human disease, remains to be confirmed.

Recently, the coding-complete BAV genome sequence (13,296 nt) was determined (Lauck et al., 2015; Walker et al., 2015), and two long-known viruses, Sweetwater Branch virus (SWBV) and Beatrice Hill virus (BHV), were identified as tibroviruses after their coding-complete genome sequences (13,141 nt and 13,296 nt, respectively) were determined (Walker et al., 2015; Huang et al., 2016; Wiley et al., 2017). SWBV was originally isolated in 1981 from biting midges (*Culicoides insignis*) together with BAV in Florida, USA (Gibbs et al., 1989). Anti-SWBV antibodies have been detected in healthy Trinidad water buffaloes (Calisher et al., 1993). BHV was first reported in 1984 as a novel virus of biting midges (*Culicoides peregrinus*) that were collected at Beatrice Hill, Northern Territory, Australia (Standfast et al., 1984).

Beyond sequencing, very little research has been performed on any tibrovirus. Results of recent studies revealed structural elements of the BASV U1 protein (Buchko et al., 2017) and indicated that tibrovirus *U3* encodes a small viroporin-like transmembrane protein (Walker et al., 2015). In addition, it is suspected that BASV *G* encodes a class III fusion glycoprotein (G) (Steffen et al., 2013). BASV G mediates pseudotyped vesicular stomatitis Indiana virus (VSIV) particle entry into a variety of human and nonhuman cells in a pH-dependent manner (Steffen et al., 2013). However, whether such particle cell entry (susceptibility) can be correlated to actual BASV replication (permissiveness) in the same cells remains unclear.

To further understand which cell types tibrovirus particles may enter and in which cell types authentic tibroviruses may replicate, we exposed 53 human, 11 bat, 7 nonhuman primate, 1 hispid cotton rat, 1 boa constrictor, and 1 Asian tiger mosquito cell lines to newly established infectious recombinant vesicular stomatitis Indiana viruses (rVSIVs) encoding the eight diverse tibrovirus Gs and to four authentic tibroviruses (BAV, CPV, SWBV, and TIBV). The obtained rVSIV infections rates, which reflect tibrovirus particle cell entry (host cell susceptibility), indicate that particles of all tibroviruses, not only those of BASV, easily enter a broad range of human and non-human cells. We confirm pH dependency of BASV G-mediated entry, show that tibrovirus particle cell entry is likely dynamin-dependent and cholesterol-independent, and extrapolate these findings to all tibroviruses. Using the four authentic tibroviruses and cross-reacting anti-tibrovirus N antibodies in western blots, we demonstrate, however, that many cell lines that are susceptible to tibrovirus G-mediated particle entry are not likely to be permissive to replication of most tibroviruses after particle entry.

## MATERIALS AND METHODS

### Viruses and virus infections

Infectious recombinant vesicular stomatitis Indiana virus expressing enhanced green fluorescent protein (eGFP) and its native glycoprotein (G) served as a control for all cell susceptibility experiments (rVSIV-VSIV G control; Figure 1A, top). Open reading frames encoding Bas-Congo virus (BASV; GenBank # JX297815.1), Ekpoma virus 1 (EKV-1; GenBank #KP324827), Ekpoma virus 2 (EKV-2; GenBank #KP324828), and Sweetwater Branch virus (SWBV; Table 1) G were synthesized by DNA2.0 (Newark, CA, USA) and cloned into the eGFP-expressing rVSIV control backbone in place of native VSIV G, as described before (Wong et al., 2010; Kleinfelter et al., 2015), yielding rVSIV-BASV G, rVSIV-EKV-1 G, rVSIV-EKV-2 G, and rVSIV-SWBV G, respectively (Figure 1A). Open reading frames encoding Bivens Arm virus (BAV), Beatrice Hill virus (BHV), Coastal Plains virus (CPV), and Tibrogargan virus (TIBV) were amplified by RT-PCR from infected cells (BAV, CPV, TIBV) or infected laboratory mouse brain samples (BHV) obtained from the World Reference Center for Emerging Viruses and Arboviruses (WRCEVA) and cloned into the rVSIV control backbone in place of native VSIV G (Wong et al., 2010; Kleinfelter et al., 2015) to yield rVSIV-BAV G, rVSIV-BHV G, rVSIV-CPV G, and rVSIV-TIBV G, respectively (Figure 1A). The sequences of the G-encoding regions of all created rVISV genomes were confirmed to be identical to those of the respective GenBank accession numbers by Sanger sequencing. Primers were designed to amplify rVSIV-tibrovirus G-encoding regions by RT-PCR, and these fragments were sequenced by Macrogen, Rockville, MD, USA. All rVSIVs were rescued, grown, and plaque-purified as described before (Wong et al., 2010; Kleinfelter et al., 2015). For rVSIV expansion, adult grivet (*Chlorocebus aethiops*) kidney Vero cells (American Type Culture Collection [ATCC], Manassas, VA, USA; #CCL-81) were exposed to rVSIVs (MOI = 0.01) and cell culture supernatants were harvested once eGFP expression was observed in ≈60-70% of the cells. Supernatants were centrifuged to remove gross debris, used to quantify virus by plaque assay on Vero cells, aliquoted, and stored at -80°C until use. Using specifically designed primers, G-encoding regions of the genomes of the expanded four authentic tibroviruses were sequenced by Macrogen to ensure that the deduced G amino acid sequences are identical to the respective deduced G sequences from tibrovirus genomes deposited to GenBank.

**Figure 1.**
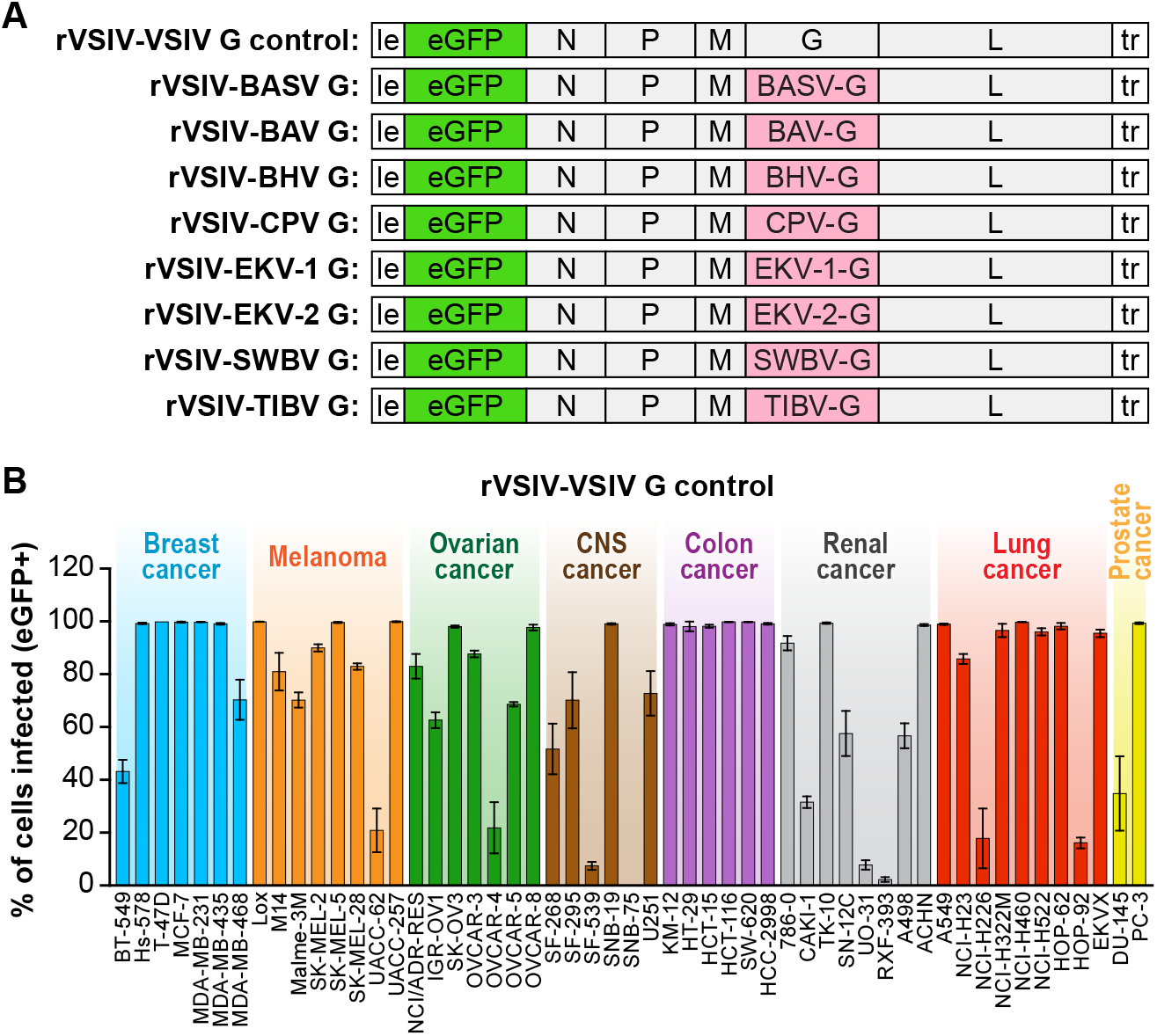
Recombinant vesiculoviruses used in this study. **(A)** Genome schematic of recombinant vesicular stomatitis Indiana virus (rVSIV) expressing its native glycoprotein (G) and enhanced green fluorescent protein (eGFP) (rVSIV-VSIV G control; top row) and rVSIVs created for this study encoding tibrovirus G instead of VSIV G (other rows). **(B)** Infectivity of rVSIV-VSIV G control (MOI = 3). The percentage of eGFP-expressing NCI-60 human cell panel cell lines was measured by high-content imaging at 24 h post-exposure. All experiments were performed in triplicate; error bars show standard deviations. BHV, Beatrice Hill virus; BASV, Bas-Congo virus; BAV, Bivens Arm virus; CNS, central nervous system, CPV; Coastal Plains virus; EKV-1, Ekpoma virus 1; EKV-2, Ekpoma virus 2; MOI, multiplicity of infection; NCI, National Cancer Institute; SWBV, Sweetwater Branch virus; TIBV, Tibrogargan virus. NCI-60 cell lines are listed by their abbreviations and grouped by organ/cancer type.

**Table 1.**
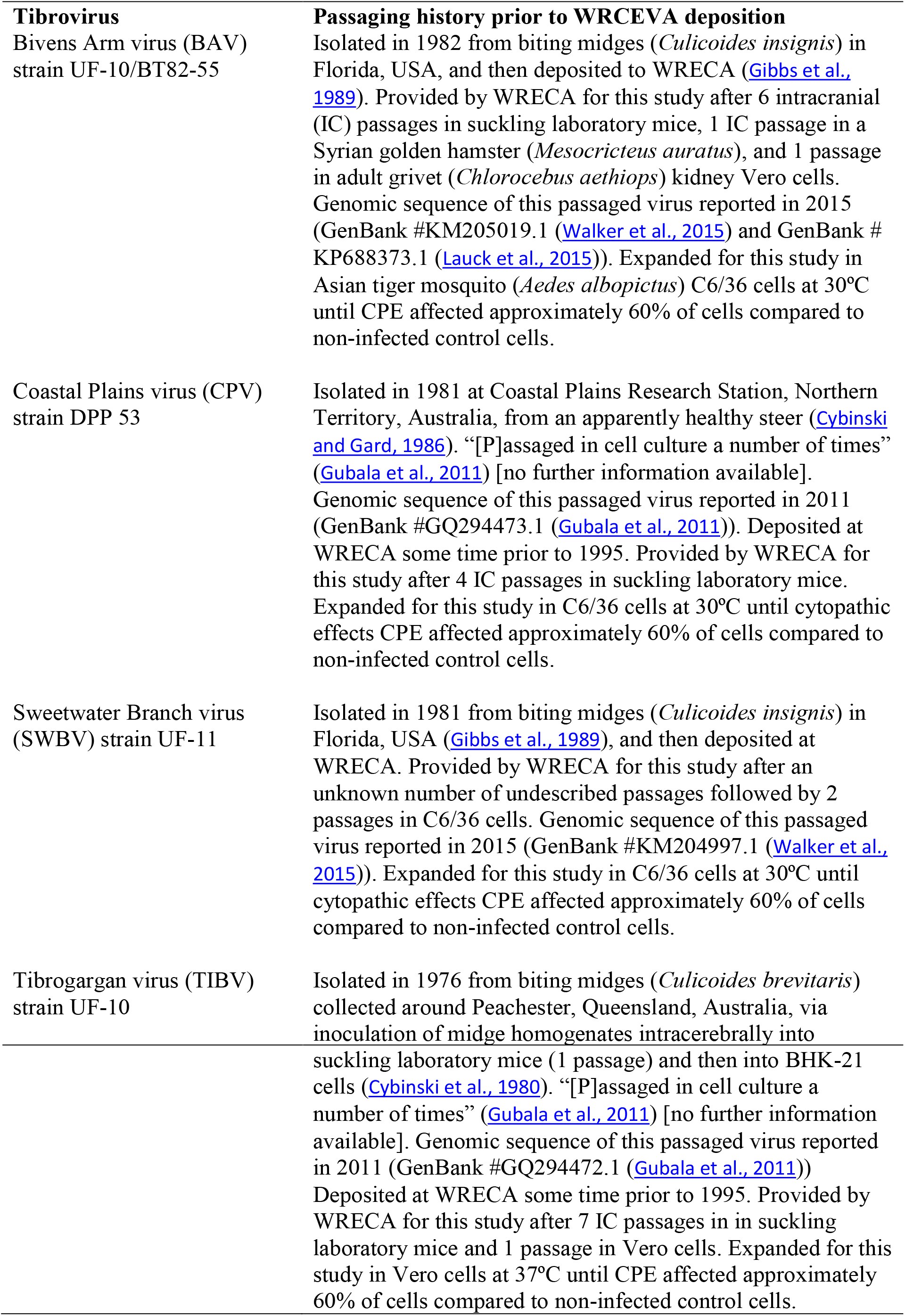
Strain and passaging history of the four authentic tibroviruses studied from the World Reference Center for Emerging Viruses and Arboviruses (WRCEVA) (The University of Texas Medical Branch (UTMB), 2018).

Information on the four authentic tibroviruses used in this study, including their sometimes incompletely documented passaging histories, is outlined in Table 1. All four viruses were received as lyophilized powders in individual tubes. Each powder was resuspended in 1 ml of phosphate-buffer saline (PBS). 100 μl of a given suspension were added to a T75 flask containing Vero or Asian tiger mosquito (*Aedes albopictus*) larva C6/36 (ATCC, #CRL-1660) cells (see below for culture conditions). Cell supernatants were harvested when cytopathic effects (CPE) affected approximately 60% of cells (designated P1; 15 ml/flask). 1 ml of P1 was used to infect a T175 flask of Vero or C6/36 cells. Supernatants were harvested when CPE affected approximately 60% of cells (designated P2; 30 ml/flask). P2 was centrifuged to remove gross debris, aliquoted, and stored at -80°C until use for all tibrovirus infections performed for this study. In addition, Beatrice Hill virus (BHV) strain Commonwealth Scientific and Industrial Research Organisation (CSIRO) 25 (GenBank #KY073493.1 (Wiley et al., 2017)) was kindly provided by Kim Blasdell, Australian Animal Health Laboratory (AAHL)/CSIRO, Geelong, Victoria, Australia. Unfortunately, attempts to grow BHV in any culture failed and therefore authentic BHV had to be omitted from this study.

### Cells lines

#### Human

The “NCI-60 panel”, a panel of 60 highly characterized human breast, central nervous system (CNS), colon, lung, melanoma, ovarian renal cancer, and prostate cancer cell lines (Weinstein, 2006), was obtained from the US National Cancer Institute’s Developmental Therapeutics Program (NCI DTP), Fort Detrick, Frederick, MD, USA. Of these 60 cells lines, 53 adherent cell lines were used for this study. In relevant figures and tables, these cell lines are grouped by tumor classification using different colors. All NCI-60 panel cell lines were grown at 37°C in a humidified 5% CO2 atmosphere in Rosswell Park Memorial Institute 1640 medium (RPMI-1640, ThermoFisher Scientific, Waltham, MA, USA) supplemented with 10% heat-inactivated fetal bovine serum (FBS, Sigma-Aldrich, St. Louis, MO, USA).

#### Bat

Eastern pipistrelle (*Pipistrellus subflavus*) adult lung PESU-B5L cells (Huynh et al., 2012) were kindly provide by Eric F. Donaldson, University of North Carolina, Chapel Hill, NC, USA. Egyptian rousette (*Rousettus aegyptiacus*) embryo Ro5T and Ro6E cells (Jordan et al., 2009) were kindly provided by Ingo Jordan, ProBioGen AG, Berlin, Germany. African straw-colored fruit bat (*Eidolon helvum*) adult kidney EidNi/41.3 cells (Biesold et al., 2011), Büttikofer’s epauletted fruit bat (*Epomops buettikoferi*) adult kidney EpoNi/22.1 cells, Daubenton's myotis (*Myotis daubentonii*) adult lung MyDauLu/47.1 cells, Egyptian rousette adult kidney RoNi7.1, hammer-headed fruit bat (*Hypsignathus monstrosus*) fetal lung HypLu/45.1 cells, hammer-headed fruit bat fetal kidney HypNi/1.1 cells (Kühl et al., 2011), and Egyptian rousette adult kidney RoNi/7.2 cells (Hoffmann et al., 2013) were kindly provided by Marcel A. Müller and Christian Drosten, Charité—Universitätsmedizin Berlin, Germany (cell were generated with funds from the German Research Council [DR 772/10-2]). Brazilian free-tailed bat (*Tadarida brasiliensis*) adult lung Tb1 Lu cells were obtained from ATCC (#CCL-88). Ro5T, Ro6E, and HypNi/1.1 cells were grown in Dulbecco's modified Eagle's medium (DMEM)/F-12 (Lonza, Walkersville, MD, USA) supplemented with 10% heat-inactivated FBS. All other bat cell lines were maintained in DMEM supplemented with 10% heat-inactivated FBS. All cells were incubated at 37°C in a humidified 5% CO2 atmosphere.

#### Nonhuman primate

Vero cells and embryonic grivet kidney MA104 cells (ATCC, #CCL-2378) were grown at 37°C in a humidified 5% CO2 atmosphere in DMEM. Primary gorilla (*Gorilla gorilla*) RpGor53, common chimpanzee (*Pan troglodytes*) S008397, and common chimpanzee RP00226 cells were obtained from Coriell Institute for Medical Research (Camden, NJ, USA) and grown under the same conditions in Eagle's Minimum Essential Medium (EMEM; Lonza) supplemented with 10% heat-inactivated FBS.

#### Rodent

Hispid cotton rat (*Sigmodon hispidus*) lung CRL cells (ATCC, #PTA-3920) were grown at 37°C in EMEM supplemented with 10% heat-inactivated FBS.

#### Reptile

Boa constrictor (*Boa constrictor*) kidney JK cells (Stenglein et al., 2012) were kindly provided by Joseph L. DeRisi, University of San Francisco, CA, USA, and grown at 30°C in a humidified 5% CO2 atmosphere in EMEM with Hanks’ balanced salt solution (HBSS, Lonza) supplemented with 10% heat-inactivated FBS.

#### Insect

Asian tiger mosquito C6/36 cells were grown at 30°C in a humidified 5% CO2 atmosphere in EMEM.

### Virus infections

Inoculation of cell lines with rVSIV-VSIV G control and rVSIVs expressing tibrovirus Gs were performed uniformly. Cell media were removed, and cells were washed once with the appropriate media without FBS. Cells were then exposed to virus particles at the indicated multiplicities of infection (MOI) and appropriate temperatures in a humidified 5% CO2 atmosphere for 1 h under gentle rocking every 15 min. Virus inocula were then removed, cells were washed once with appropriate media without FBS, and then incubated in the appropriate growth media containing 2% heat-inactivated FBS at the appropriate temperatures in a humidified 5% CO2 atmosphere for the indicated times. At 24 h later, cells were fixed in 10% neutral buffered formalin (ThermoFisher Scientific), and the total numbers of cells were counted using cell membrane-permeant, minor groove-binding blue fluorescent Hoechst 33342 DNA stain (Cell Signaling Technology, Danvers, MA, USA). The authentic tibroviruses used in this study did not cause plaques on any tested cell line and qRT-PCR assays have not yet been established for any of them. Hence, equal volumes of virus powder suspensions were used to expose identical numbers of cells. Cells were exposed to authentic tiborviruses in a humidified 5% CO2 atmosphere.

### Inhibitor studies

Ammonium chloride, chloroquine, chlorpromazine, concanamycin A, and dynasore were obtained from Sigma-Aldrich. Dyngo-4a was kindly provided by Adam McCluskey and Volker Haucke, Freie Universität Berlin, Germany. Confluent Vero cells were exposed to the indicated concentrations of inhibitors in the appropriate medium for 30 min. rVSIV-VSIV G control or rVSIV expressing tibrovirus Gs were directly added to inhibitor-containing media. Cells were exposed for 1 h at 37°C (MOI = 0.6), followed by eGFP quantification 16 h later using an Infinite M1000 microplate reader (Tecan, Männedorf, Switzerland). The Cell Counting Kit-8 (Dojindo Molecular Technologies, Rockville, MD, USA) was used according to the manufacturer’s instructions to determine the cytotoxicity of inhibitors in uninfected cells in parallel to infection assays.

### Detection of tibrovirus infection

Infection by rVSIV-VSIV G control and rVSIVs expressing tibrovirus glycoproteins was measured by detecting eGFP expression using the Operetta High-Content Imaging System with Harmony 3.1 analysis software (PerkinElmer, Shelton, CT, USA).

Anti-CPV N and anti-TIBV N antibodies were custom-made by ThermoFisher Scientific. Due to the large scale of the study, and because these antibodies were unfortunately not suitable for indirect fluorescence assays (IFA), authentic tibrovirus infection was determined via detection of tibrovirus nucleoprotein (N) expression by western blot. Briefly, CPV and TIBV N peptides were designed using Antigen Profiler software (ThermoFisher Scientific). Two 19-mers were designed, produced, and injected into rabbits by the company. Rabbit sera containing antibodies were harvested and purified by the company 72 days after peptide injection. To detect authentic tibroviruses, virus-exposed cells were harvested and lysed using Cell Lysis Buffer (Cell Signaling Technology) with cOmplete Protease Inhibitor Cocktail (Sigma) (once cytopathic effects [CPE] appeared—otherwise at day 10 post-infection; Tables 2 and 3). Protein concentrations in each sample were measured using the bicinchoninic acid (BCA) assay (ThermoFisher Scientific) following the manufacturer’s instructions. Lysates were then analyzed by western blots. Protein samples (20 μg) were loaded into the wells of NuPAGE 4-12% Bis-Tris Gels (ThermoFisher Scientific), and gels were run with MOPS SDS Running Buffer (ThermoFisher Scientific). Gels were dry-transferred using the iBlot 2 Gel Transfer Device (ThermoFisher Scientific). Primary antibodies were diluted to 1:1,000; goat anti-rabbit IgG coupled to horseradish peroxidase (HRP) (Sigma Aldrich) was diluted to 1:3,000. Protein loading was controlled by detecting cellular β-actin using anti-beta actin antibody ab8227 (Abcam, Cambridge, MA, USA). Western blotting was performed using a BlotCycler (Biocompare, South San Francisco, CA, USA) after transfer. Tibrovirus N bands were detected with SuperSignal West Femto Maximum Sensitivity Substrate (ThermoFisher Scientific). Gel images were taken using a G:BOX gel documentation system (Syngene, Frederick, MD, USA).

**Table 2.**
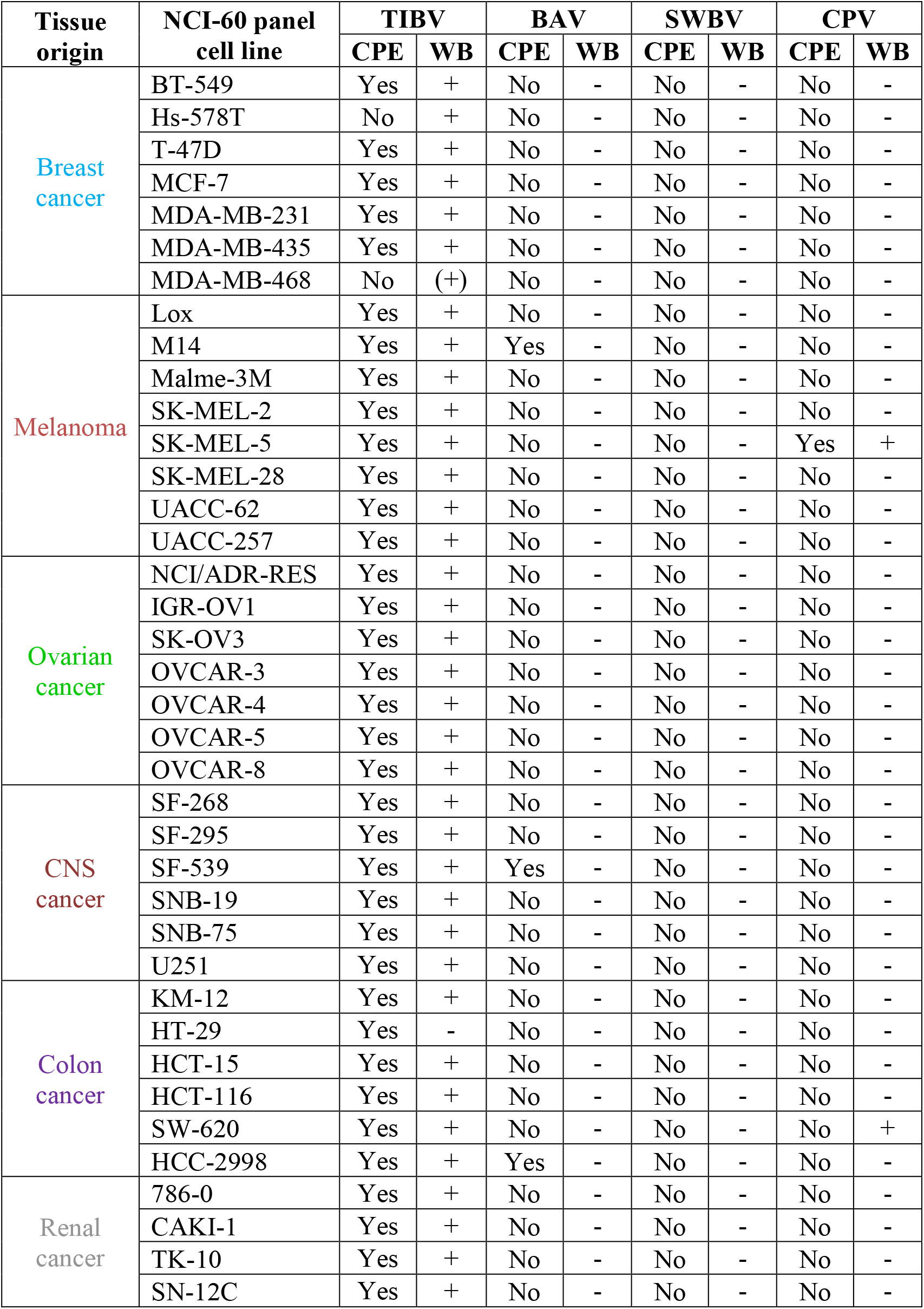
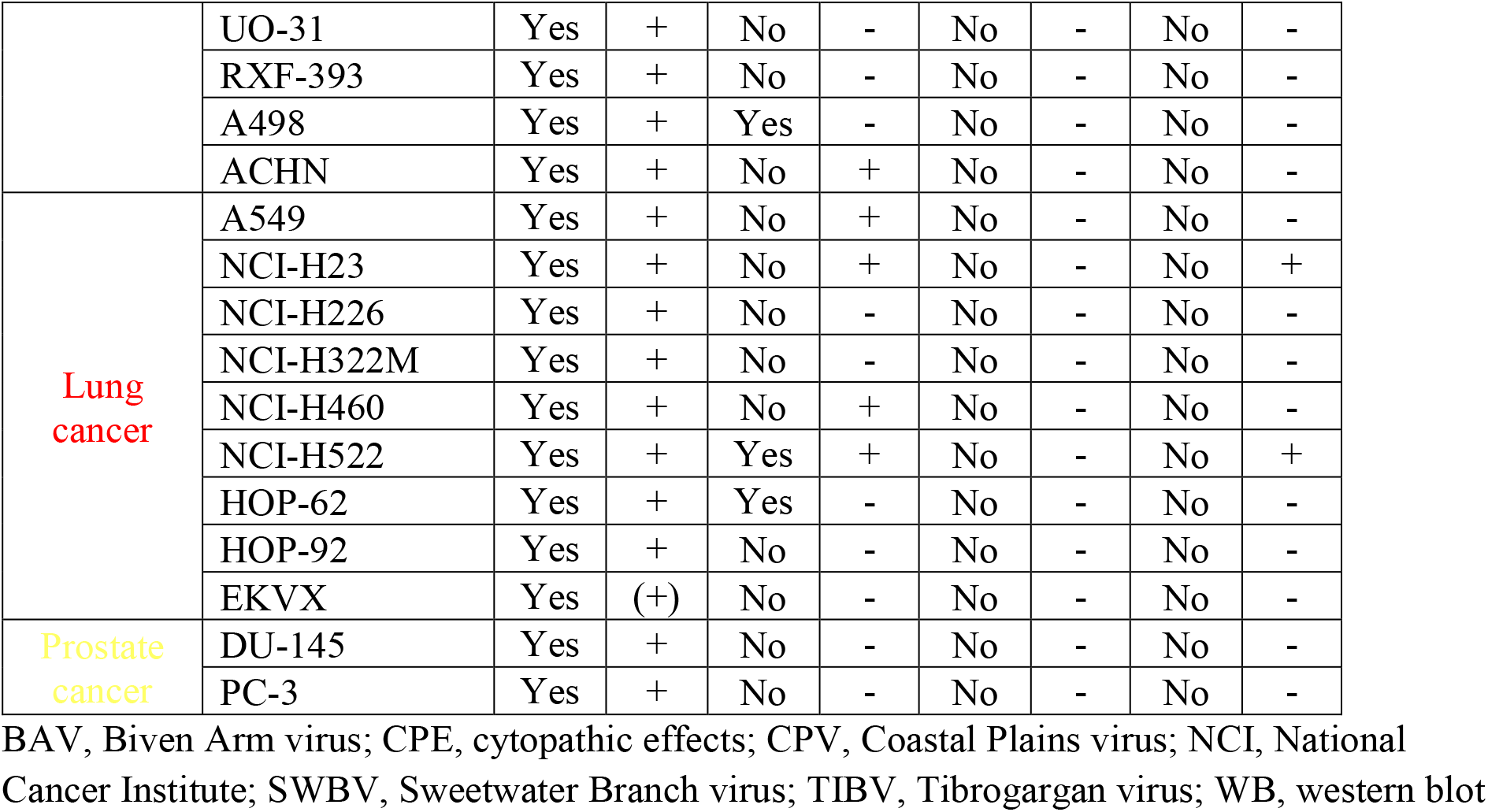
Detection of authentic tibroviruses in human cell lines.

**Table 3.**
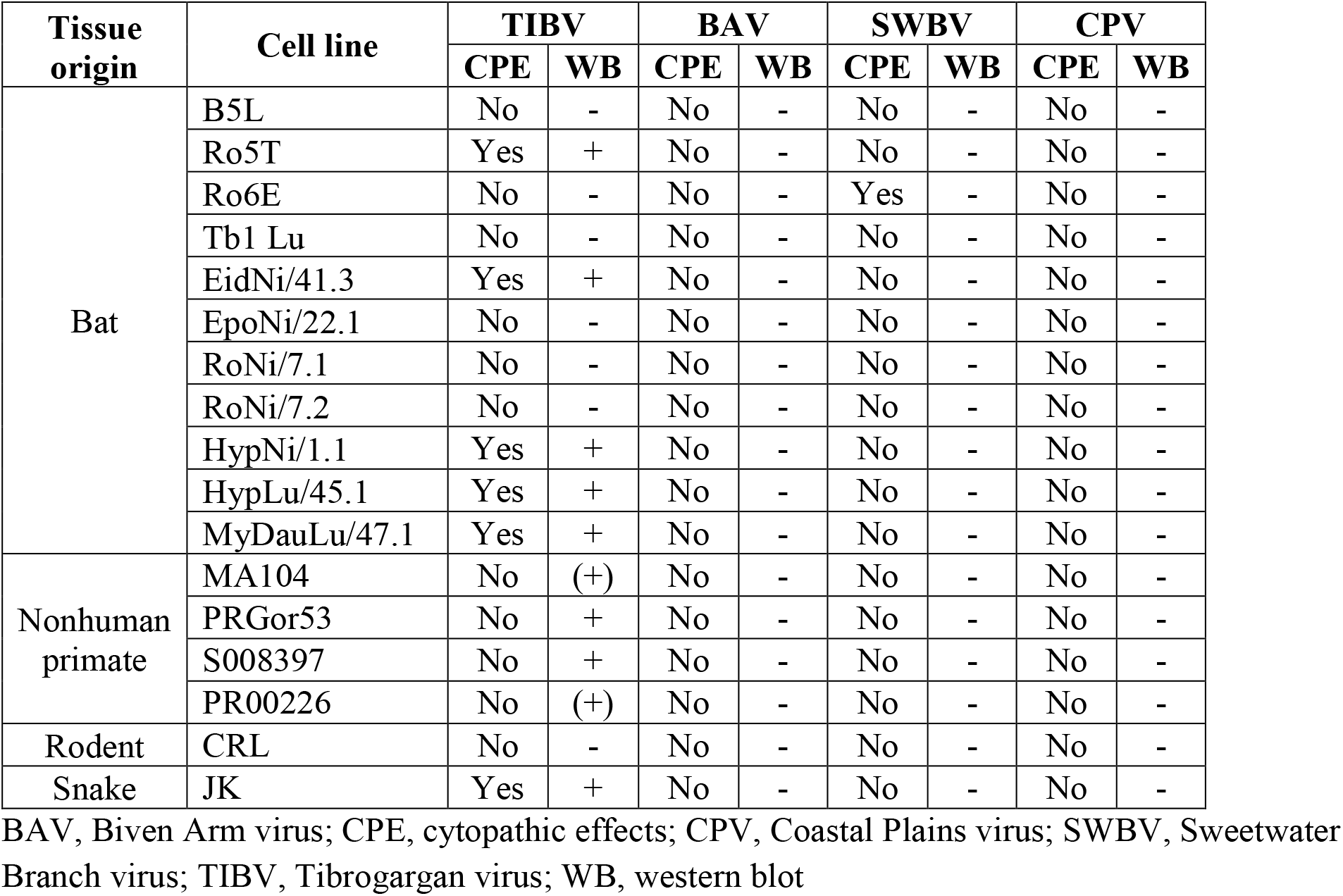
Detection of authentic tibroviruses in nonhuman cell lines.

## RESULTS

### Tibrovirus glycoproteins mediate virion entry into a broad range of human and animal cell types

The glycoprotein (G) is the only rhabdovirus genome-encoded protein determining rhabdovirion cell entry (Altstiel and Landsberger, 1981; Wunner et al., 1984; Gaudin et al., 1993; Regan and Whittaker, 2013). Cell entry is thought to occur after G engages virus-specific cell-surface receptors or attachment factors (Bearzotti et al., 1999; Lafon, 2005; Finkelshtein et al., 2013) and subsequently mediates low-pH-induced viral and endosomal membrane fusion (Florkiewicz and Rose, 1984; Blumenthal et al., 1987; Sun et al., 2005). Vesicular stomatitis Indiana virus (VSIV; *Rhabdoviridae: Vesiculovirus: Indiana vesiculovirus*) has an extremely broad, close-to-universal, cellular tropism. This broad tropism is due the fact that most cells are susceptible to VSIV G-mediated virus particle entry (cells allow virion entry) and that most cells are also permissive to VSIV replication after particle entry (cells do not restrict virus replication or virion egress) (Hastie et al., 2013). Using reverse genetics to create recombinant VSIV (rVSIV), VSIV G can easily be replaced with other mononegaviral glycoproteins, thereby possibly changing the receptor engagement of the rVSIV to that of G of the heterotypic virus, while likely maintaining the ability of the recombinant virus to replicate in most cells after particle entry (change of cell susceptibility while maintaining the same cell permissiveness). Therefore, rVSIVs expressing heterotypic Gs can be used to perform well-controlled, initial cell susceptibility evaluations of G from heterotypic viruses without including the authentic heterotypic viruses in the experiment (Garbutt et al., 2004; Tani et al., 2011; Hastie et al., 2013; Moreira et al., 2016; Robinson and Whelan, 2016; Raaben et al., 2017; Jangra et al., 2018). This approach is especially useful when the actual heterotypic viruses are only known from sequences, i.e., have not been isolated in culture (e.g., Bas-Congo virus [BASV], Ekpoma virus 1 [EKV-1], Ekpoma virus 2 [EKV-2]).

To improve understanding of tibrovirus cell tropism, and to assess whether *in vitro* cell tropism of tibroviruses might be indicative of (human) host tropism, we used a wild-type rVSIV expressing its native VSIV G and enhanced green fluorescent protein (eGFP) (rVSIV-VSIV G control), and created rVSIVs based on this control expressing the Gs of all eight known tibroviruses (BASV, Beatrice Hill virus [BHV], Bivens Arm virus [BAV], Coastal Plains virus [CPV], EKV-1, EKV-2, Sweetwater Branch virus [SWBV], and Tibrogargan virus [TIBV]) in place of VSIV G (Figure 1A). We then exposed 53 well-characterized human adherent cancer cell lines of the “NCI-60 panel” (Weinstein, 2006) to rVSIV-VSIV G control and the eight rVSIVs encoding tibrovirus Gs (MOI = 3) and measured virion cell entry (cell susceptibility to tibrovirus G-mediated particle entry) by quantifying the percentage of eGFP-expressing cells using high-content imaging at 24 h post-exposure (Figures 1B and 2). As expected, rVSIV-VSIV G control infected almost all cell lines, with the notable exceptions of one central nervous system (CNS) cancer cell line (SNB-75) and one renal cancer cell line (RXF-393). Hence, these two cell lines are not susceptible, not permissive, or not susceptible and not permissive to VSIV infection. rVSIV-VSIV G control infection occurred in 100% of cells in 25 of the 53 tested cell lines, and infection efficiency was approximately ≤50% in 8 of them (Figure 1B).

**Figure 2.**
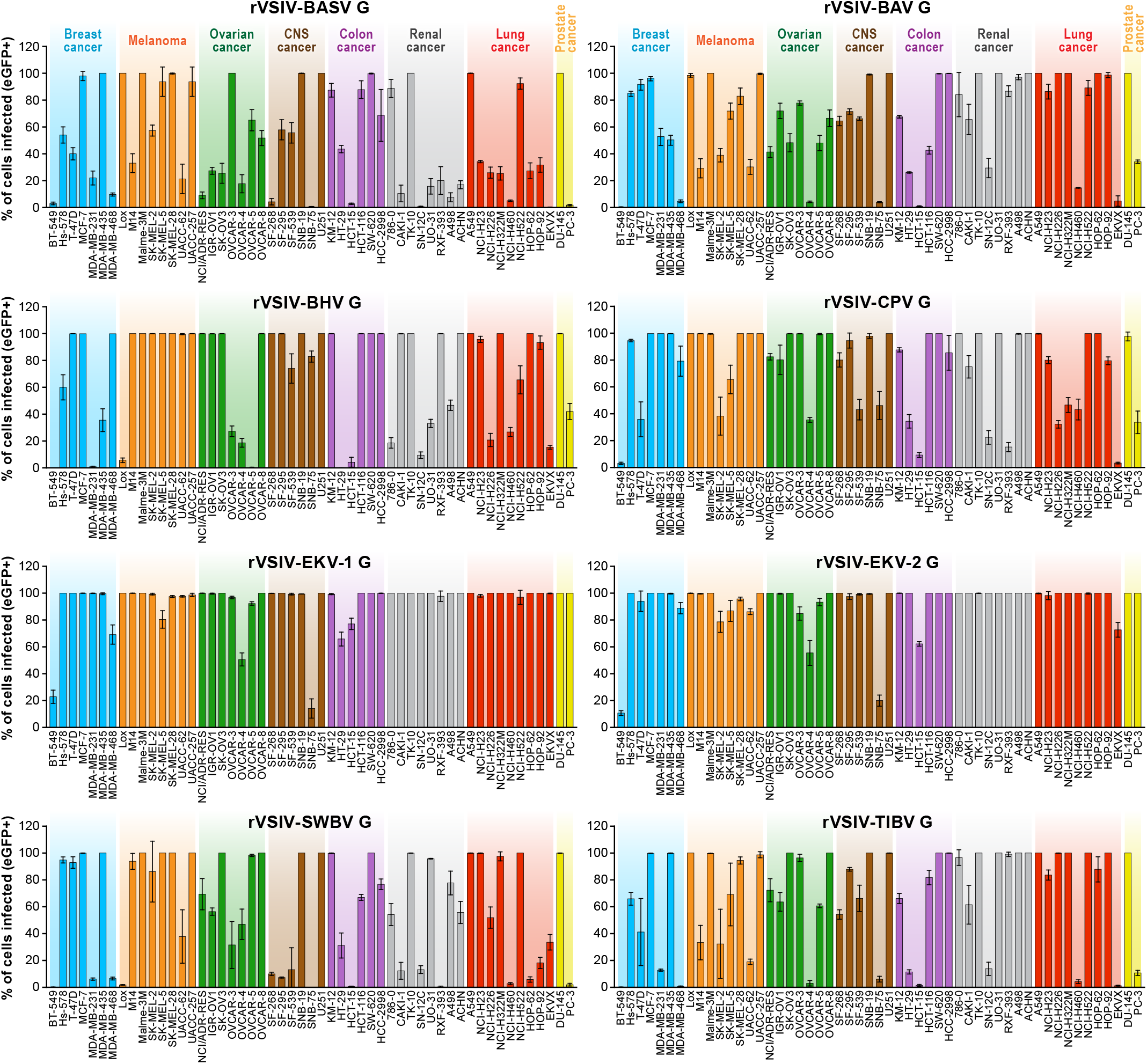
Tibrovirus glycoproteins mediate virion entry into a broad range of human cell types. Same experiment as in Figure 1B using rVSIVs expressing diverse tibrovirus G glycoproteins (MOI = 3). The percentage of eGFP-expressing NCI-60 human cell panel cell lines was measured by high-content imaging at 24 h post-exposure. All experiments were performed in triplicates; error bars show standard deviations. BHV, Beatrice Hill virus; BASV, Bas-Congo virus; BAV, Bivens Arm virus; CNS, central nervous system, CPV; Coastal Plains virus; EKV-1, Ekpoma virus 1; EKV-2, Ekpoma virus 2; eGFP, enhanced green fluorescent protein; MOI, multiplicity of infection; NCI, National Cancer Institute; SWBV, Sweetwater Branch virus; TIBV, Tibrogargan virus; rVSIV, recombinant vesicular stomatitis Indiana virus. NCI-60 cell lines are listed by their abbreviations and grouped by organ/cancer type.

Due to the overall known broad cell tropism of VSIV (Hastie et al., 2013), and assuming that VSIV uses a different cell-surface receptor or attachment factor than tibroviruses, we expected rVSIVs expressing tibrovirus Gs in place of VSIV G to enter fewer cell lines with less efficiency than rVSIV-VSIV G control. Assuming that within-cell VSIV restriction factors do not target G, we further surmised that rVSIV-VSIV G control and rVSIVs expressing a tibrovirus G would replicate at similar efficiencies in cells that permit entry of both viruses, i.e., that these cells would be equally permissive. Interestingly, rVSIV-BAV (3 negative cell lines, 19 cell lines with ≈100% infection efficiency), rVSIV-BASV (4 negative cell lines, 11 cell lines with ≈100% infection efficiency), rVSIV-SWBV (6 negative cell lines, 21 cell lines with ≈100% infection efficiency), and rVSIV-TIBV (5 negative cell lines, 23 cell lines with ≈100% infection efficiency) fulfilled the first assumption, but not necessarily the second (Figure 2). Even more surprisingly, rVSIVs expressing Gs of the remaining tibroviruses equaled or outperformed rVSIV-VSIV G control in number of cell types infected and/or infection efficiency. Most notably, rVSIV-EKV-1 and rVSIV-EKV-2 entered and replicated in all tested cell lines. Only two cell lines were infected with approximately ≤50% efficiency, and 44 cell lines (rVSIV-EKV-1) or 40 cell lines (rVSIV-EKV-2) were infected with ≈100% efficiency (Figure 2). Importantly, cell lines resistant to rVSIV-VSIV G control infection (i.e., CNS SNB-75, RXF-393) were not necessarily resistant to infection with rVSIVs expressing certain tibrovirus Gs. For instance, the SNB-75 cell line was efficiently infected with rVSIV-BHV (≈80%) and rVSIV-CPV (≈45%). The cell line was inefficiently infected with rVSIV-BAV, rVSIV-EKV-1, rVSIV-EKV-2, and rVSIV-TIBV and not at all infected with rVSIV-BASV and rVSIV-SWBV. Likewise, the renal cancer RXF-393 cell line was efficiently infected with rVSIV-BAV, rVSIV-BHV, rVSIV-EKV-1, rVSIV-EKV-2, and rVSIV-TIBV, inefficiently infected with rVSIV-BASV and rVSIV-CPV, and resistant to rVSIV-SWBV. Together, these results indicate that all tibroviruses may be able to broadly enter human cell types and that individual tibrovirus Gs bestow different cell tropism patterns. Furthermore, these data indicate that resistance of SNB-75 and RXF-393 cells to rVSIV-VSIV G control occurs at the VSIV particle entry step and is not due to replication restriction as rVSIVs encoding some non-VSIV Gs were able to replicate in these cells.

To determine whether tibrovirus Gs also mediate virion entry into nonhuman cell lines, we exposed 11 bat, 5 nonhuman primate, 1 hispid cotton rat, 1 boa constrictor, and 1 Asian tiger mosquito cell lines to rVSIV-VSIV G control and the 8 rVSIVs expressing tibrovirus Gs (Figure 3). Two bat cell lines, PESU-B5L and MyDauLu/47.1, were almost uniformly resistant to infection (virus particle entry and/or subsequent replication), and only a low level of infection by rVSIV-SWBV was detected in PESU-B5L cells. Tb1 Lu cells were resistant to rVSIV-BASV, rVSIV-EKV-1, and rVSIV-VSIV G control but were susceptible to all other viruses. RoNi/7.1 cells were resistant to rVSIV-BASV, rVSIV-BHV, rVSIV-CPV, rVSIV-EKV-1, and rVISV-TIBV. All other bat cell lines were variably susceptible and permissive to all rVSIVs (Figure 3A). All 9 rVSIVs infected hispid cotton rat CRL, boa constrictor JK, and nonhuman primate MA104 and Vero cells with high efficiency. However, gorilla (RPGor53) cells could not be infected by rVSIV-VSIV G control or rVSIV-SWBV, and common chimpanzee (S008397 and RP00226) cells were highly susceptible to rVSIV-CPV but much less susceptible or resistant to the other rVSIVs expressing tibrovirus Gs (Figure 3A). Finally, all viruses infected Asian tiger mosquito (C6/36) cells, but infection only became apparent at ≈72 h post-exposure. Interestingly, rVSIV-VSIV G control infected these cells least efficiently compared to all other viruses (Figure 3B). Together, these results indicate that tibroviruses may be able to broadly enter non-human cell lines, but some cell lines (e.g., PESU-B5L and MyDauLu/47.1) may not be permissive to VSIV replication after particle entry.

**Figure 3.**
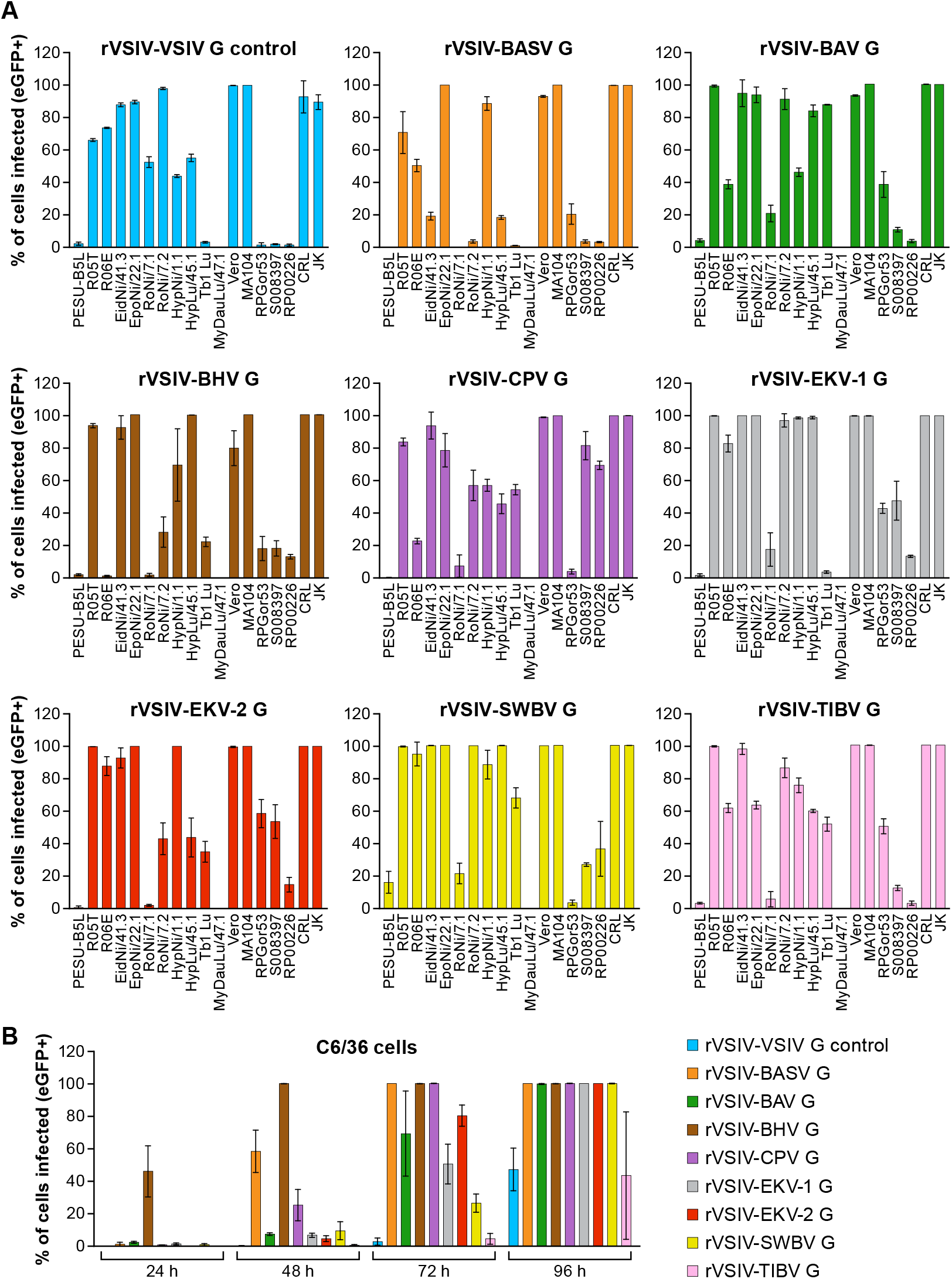
Tibrovirus glycoproteins mediate virion entry into a broad range of animal cell types. Same experiment as in Figures 1B and 2 using different cell types exposed to rVSIV-VSIV G control and rVSIVs expressing diverse tibrovirus glycoproteins (G) (MOI=3). **(A)** Bat (PESU-B5L, Ro5T, Ro6E, EidNi/41.3, EpoNi/22.1, RoNi/7.1, RoNi/7.2, HypNi/1.1, HypLu/45.1, Tb1 Lu, MyDauLu/47.1), nonhuman primate (Vero, MA104, RPGor53, S008397, RP00226), hispid cotton rat CRL, and boa constrictor JK cell lines. **(B)** Asian tiger mosquito C6/36 cells. The percentage of eGFP-expressing cell lines was measured by high-content imaging at 24 h post-exposure (bat, nonhuman primate, hispid cotton rat, and boa constrictor cell lines) or at 24 h, 48 h, 72 h, and 96 h post-exposure (Asian tiger mosquito cells). All experiments were performed in triplicate; error bars show standard deviations. BHV, Beatrice Hill virus; BASV, Bas-Congo virus; BAV, Bivens Arm virus; CPV, Coastal Plains virus; eGFP, enhanced green fluorescent protein; EKV-1, Ekpoma virus 1; EKV-2, Ekpoma virus 2; MOI, multiplicity of infection; SWBV, Sweetwater Branch virus; TIBV, Tibrogargan virus; rVSIV, recombinant vesicular stomatitis Indiana virus.

The experiments above were performed with high amounts of rVSIVs (MOI = 3) as our primary objective was to determine whether different cells types are absolutely susceptible or resistant to tibrovirus G-mediated particle entry. To determine relative differences, we repeated the experiments with a MOI = 0.3 (Supplementary Figures 1 and 2). As expected, percentages of rVSIV-infected cells were generally decreased in the lower MOI experiment, but numerous cell lines were still infected at the 100% level. Most strikingly, rVSIV-EKV-1 and rVSIV-EKV-2 infection rates among human cell lines barely diminished, suggesting that EKV-1 and EKV-2 are more efficient in entering human cell lines compared to all other tibroviruses.

### Tibrovirus particle host cell entry is dependent on low pH and dynamin but is independent of cholesterol

Rhabdovirion cell entry is thought to generally occur by a pH-dependent mechanism (Florkiewicz and Rose, 1984; Blumenthal et al., 1987; Sun et al., 2005). Previously, rVSIV particles devoid of VSIV G and pseudotyped with BASV G were also confirmed to enter target cells in a pH-dependent manner (Steffen et al., 2013). To evaluate whether all tibrovirus Gs function analogously, we chose a common cell line universally susceptible to tibrovirus G-mediated particle entry (Vero cells, Figure 3A). These cells were pretreated with increasing concentrations of endosomal pH modulators (lysosomotropic weak bases NH4Cl or chloroquine or vacuolar type H+-ATPase inhibitor concanamycin A) for 30 min followed by exposure to rVSIV-VSIV G control and all 8 rVSIVs expressing tibrovirus Gs at MOI = 0.6 in the presence of these modulators for 1 h. At 16 h, we quantified eGFP expression. As previously described (Steffen et al., 2013), rVSIV-VSIV G control and rVSIV-BASV infectivity decreased with increasing modulator concentrations in the absence of modulator-induced cytotoxicity. As expected, similar results were obtained with all other rVSIVs (Figure 4A).

**Figure 4.**
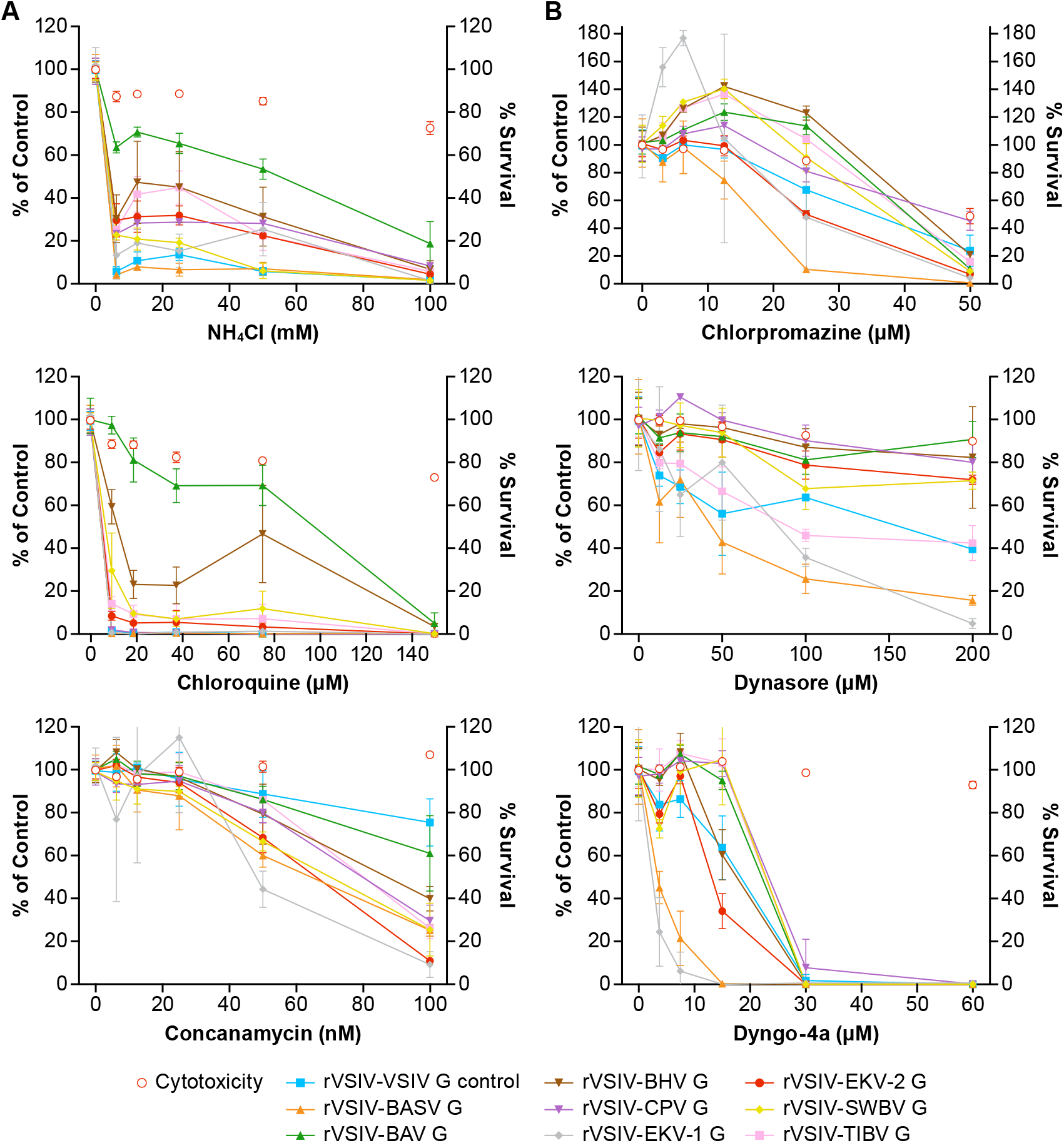
Tibrovirus particle host cell entry is dependent on low pH and dynamin but is independent of cholesterol. Tibrovirus G glycoprotein-mediated cell entry occurs via a low-pH-dependent, clathrin-mediated endocytosis-like pathway. **(A)** Effects of pretreatment of grivet (Vero) cells with increasing concentrations of endosomal pH modulators on cell entry of rVSIV-VSIV G control and rVSIVs expressing diverse tibrovirus glycoproteins (G). **(B)** Effect of pretreatment of Vero cells with increasing concentrations of clathrin-mediated endocytosis inhibitors on cell entry of the same viruses as in **A.** Cells were pretreated with the indicated concentrations of inhibitors for 30 min and then exposed to rVSIVs (MOI = 0.6) in the presence of inhibitors for 1 h at 37°C, followed by removal of virus inocula. Total expression levels of eGFP were measured using a Tecan microplate reader at 16 h post-exposure. BHV, Beatrice Hill virus; BASV, Bas-Congo virus; BAV, Bivens Arm virus; CPV, Coastal Plains virus; eGFP, enhanced green fluorescent protein; EKV-1, Ekpoma virus 1; EKV-2, Ekpoma virus 2; MOI, multiplicity of infection; SWBV, Sweetwater Branch virus; TIBV, Tibrogargan virus; rVSIV, recombinant vesicular stomatitis Indiana virus.

Rhabdovirions are thought to generally enter cells by clathrin-mediated endocytosis (CME) (Sun et al., 2005; Liu et al., 2011; Cheng et al., 2012; Piccinotti et al., 2013; Weir et al., 2014; Shao et al., 2016). To determine whether this idea can be extended to tibroviruses, we pretreated Vero cells with CME inhibitors (AP-2-clathrin-coated pit lattice interrupter chlorpromazine or dynamin inhibitors dynasore and dyngo-4a) and infected the cells as described above. As expected, infectivity of all rVSIVs was inhibited by all three compounds in a dose-dependent manner in the absence of compound-induced cytotoxicity (Figure 4B). Together, these results suggest that tibrovirus G-mediated cell entry occurs analogous to that of other rhabdoviruses by low-pH-induced CME.

### Authentic tibrovirion cell entry and infections

Finally, we evaluated whether tibrovirus cell tropism data obtained with rVSIVs reflect authentic tibrovirus infections. For this experiment, we obtained samples containing the five tibroviruses that were previously isolated in culture (BAV, BHV, CPV, SWBV, and TIBV) (Cybinski et al., 1980; Standfast et al., 1984; Cybinski and Gard, 1986; Gibbs et al., 1989). We succeeded in cultivating all of these viruses except BHV and therefore continued all experiments with the remaining four viruses. Cultured isolates of BASV, EKV-1, and EKV-2 are not available and, therefore, could not be included in the experiment.

The large size of the experiment and the absence of commercially available or generally established tibrovirus detection methods were challenging. Because the first indication of ongoing rhabdovirus replication is the production of the rhabdovirus nucleoprotein (N) (Abraham and Banerjee, 1976; Ball and White, 1976; Iverson and Rose, 1981), we aimed to detect tibrovirus cell infection using anti-tibrovirus N antibodies via western blotting using newly created anti-CPV N and anti-TIBV N antibodies. First, these antibodies were tested for cross-reactivity against all four tibroviruses in exposed Asian tiger mosquito and Vero cells. As expected, anti-TIBV N antibody detected TIBV in these TIBV-infected cells. Anti-CPV N antibody detected CPV in CPV-exposed Asian tiger mosquito cells but not in Vero cells. Encouragingly, anti-TIBV N antibody also detected BAV (but not CPV and only weakly SWBV), and anti-CPV N antibody also detected SWBV (but not TIBV and only very weakly BAV) (Figure 5A).

**Figure 5.**
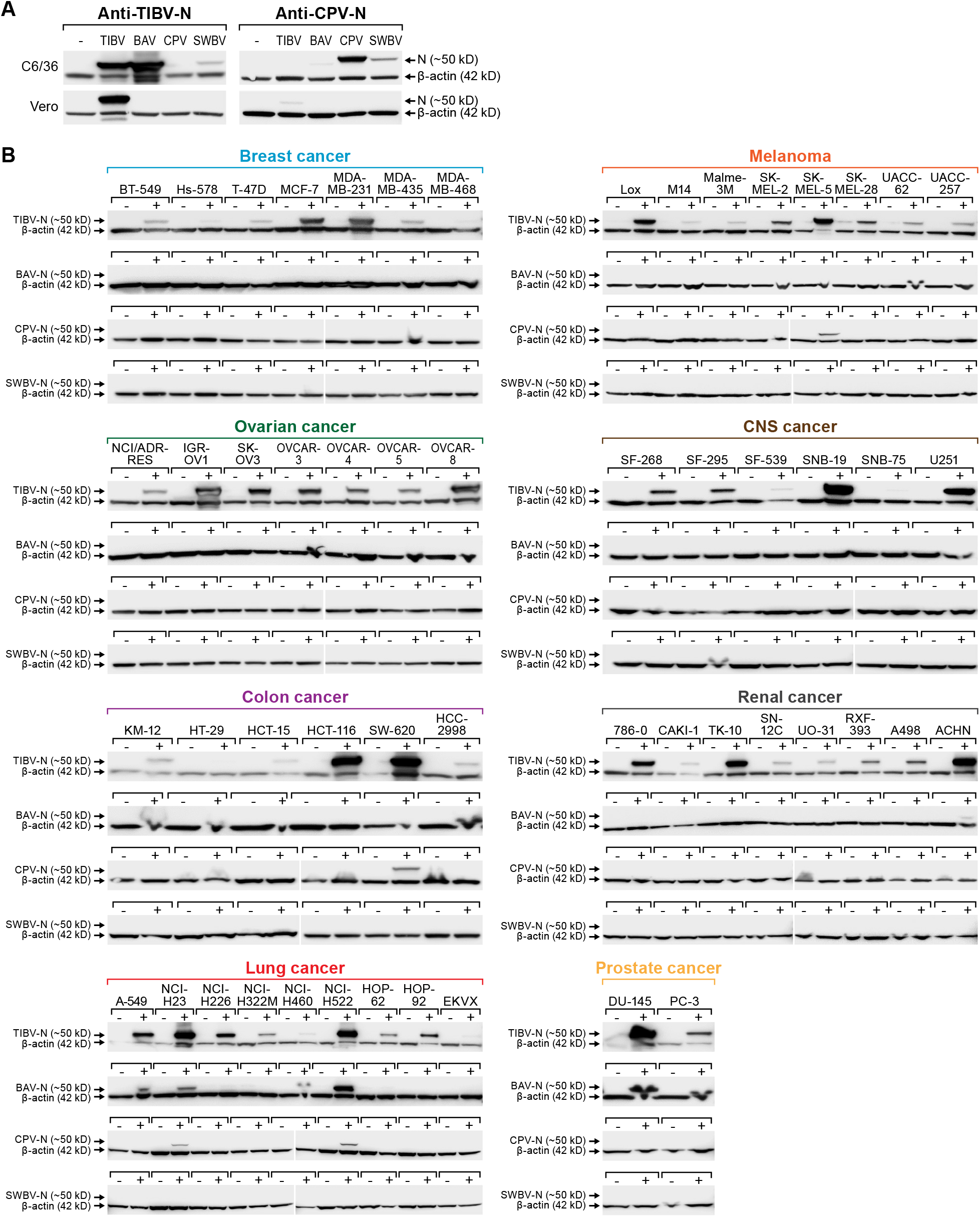
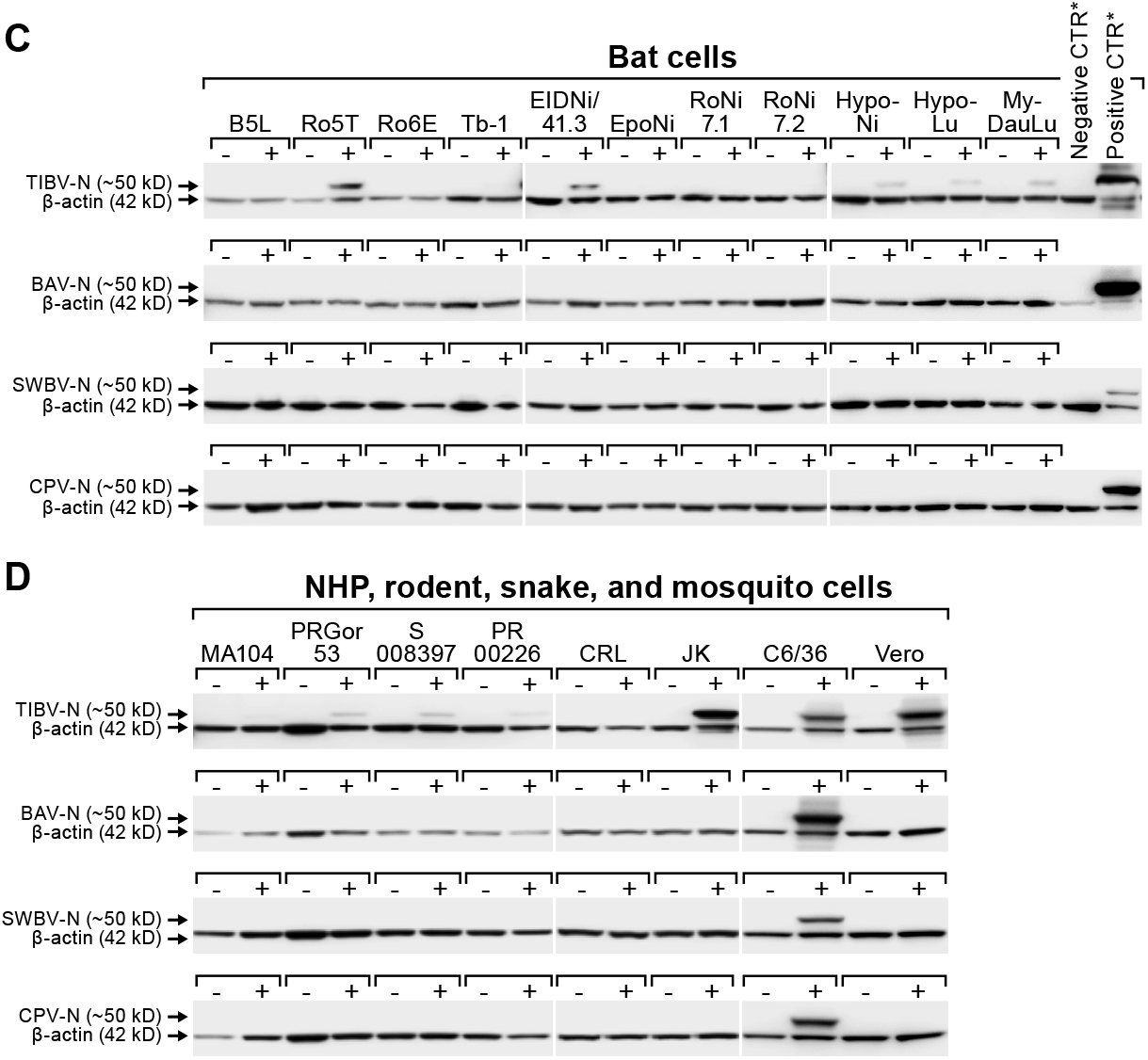
Authentic tibrovirion cell entry and infections. **(A)** Asian tiger mosquito (C6/36) and grivet (Vero) cells were exposed to medium-only control (-), BAV, CPV, SWBV, or TIBV particle preparations. Virion entry was detected via western blotting using an anti-TIBV N antibody (here shown to be strongly cross-reactive with BAV in C6/36 cells and weakly crossreactive with SWBV) and an anti-CPV N antibody (here shown to be cross-reactive with SWBV in C6/36 cells and very weakly cross-reactive with BAV). **(B)** NCI-60 cells, **(C)** bat cells, and (D) nonhuman primate (NHP), hispid cotton rat, boa constrictor, and Asian tiger mosquito cells (see also Figure 3) were exposed to medium-only control (-) or BAV, CPV, SWBV, or TIBV (+). Cell lysates were harvested after the appearance of cytopathic effects (CPE) or otherwise at day 10 post-exposure (Table 2). Tibovirion entry was detected via western blotting using the appropriate anti-TIBV, BAV, SWBV, and anti-CPV N antibodies. Protein loading was controlled by detecting ß-actin. BAV, Bivens Arm virus; CPV, Coastal Plains virus; N, nucleoprotein; NCI, National Cancer Institute; SWBV, Sweetwater Branch virus; TIBV, Tibrogargan virus.

We next exposed all human, bat, nonhuman primate, hispid cotton rat, boa constrictor, and Asian tiger mosquito cell lines used in previous experiments (Figures 1–3) to the four authentic tibroviruses and used the respective antibodies to detect N via western blotting once CPE appeared or otherwise at day 10 post-infection (Tables 2 and 3). In combination with the results depicted in Figures 2 and 3, the results of this experiment suggest that authentic tibrovirus replication does not occur in most cell types after particle entry. For instance, BAV N could be detected in only six cell lines (4 human lung cancer cell lines, 1 human prostate cancer cell line, and Asian tiger mosquito cells) (Figure 5B and 5D, Table 2), whereas rVSIV-BAV infected 50 human cell lines (Figure 2) and all other cell lines with the exception of one bat cell line (MyDauLu/47.1) (Figure 3, Table 3). Likewise, CPV infected only 4 human cell lines and Asian tiger mosquito cells, and SWBV did not infect any cell line but Asian tiger mosquito cells (Figure 5B and 5D, Tables 2 and 3). The exception to these observations was TIBV, the tibrovirus with the broadest tropism. TIBV N could be detected in the majority of tested cell lines (Figure 5B–D, Tables 2 and 3), which was largely in agreement with rVSIV-TIBV data (Figures 2 and 3). The N detection results were also largely in agreement with the observation of CPE. For instance, TIBV caused CPE in almost all human cell lines but N could be detected in two cell lines (human breast cancer Hs-578T and MDA-MB-468) in the absence of CPE (Table 2). In nonhuman cell lines, N was undetectable in cell lines that did not develop CPE with the exception of four non-human primate cell lines, which expressed N despite the absence of CPE (Table 3). In contrast, CPE and N detection was largely absent in all cell lines infected with BAV, CPV, or SWBV with some individual exceptions. Generally, N could not be detected in cell lines that did not develop CPE and N could be detected in cells that did develop CPE. However, in some cases CPE developed despite the absence of N detection and vice versa (Tables 2 and 3).

## DISCUSSION

Tibroviruses are highly undercharacterized rhabdoviruses with unknown pathogenic potential. All isolated tibroviruses (BAV, BHV, CPV, SWBV, and TIBV) have only been found in biting midge vectors or in various, apparently healthy non-human mammals (Cybinski et al., 1980; Standfast et al., 1984; Cybinski and Gard, 1986; Gibbs et al., 1989). The discovery of BASV, EKV-1, and EKV-2 genomes in human sera (Grard et al., 2012; Stremlau et al., 2015) suggests that at least some tibroviruses could infect humans. Because BASV was discovered in the serum of a severely ill individual, tibroviruses should be considered potential human pathogens for surveillance purposes until this hypothesis is ruled out.

In this initial study, we aimed to comprehensively characterize the host and cell type tropism of tibroviruses to determine whether tibroviruses in general can enter human and other animal cells. Previous studies on this topic have been extremely limited. BAV and SWBV were isolated in Asian tiger mosquito (C6/36) cells, and all attempts to grow them in baby golden hamster (*Mesocricetus auratus*) kidney (BHK-21) and adult grivet (*Chlorocebus aethiops*) kidney (Vero) cells failed (Gibbs et al., 1989). BHV was isolated only in porcine stable-equine kidney (PS-EK) cells (Standfast et al., 1984) and in C6/36 cells after a single Vero cell passage (Huang et al., 2016). CPV was isolated in BHK-21, C6/36 cells, and Vero cells (Cybinski and Gard, 1986). TIBV was originally isolated in suckling laboratory mice and could be grown in BHK-21 and Vero cells (Cybinski et al., 1980). To our knowledge, no further data on cell or animal tropism of replicating tibroviruses have been reported. However, in 2013, Steffen *et al*. reported the evaluation of BASV cell entry using an eGFP-expressing VSIV pseudotyped with BASV G (Steffen et al., 2013). Using this system, the authors demonstrated that BASV G successfully mediates VSIV particle entry into human B lymphocytes (B-THP), cervix (HeLa), colon (SW480), and colon carcinoma (CaCo-2) cells, erythroblasts (HEL), fibrosarcoma (HT1080), liver (Huh-7.5), and lung (A549) cells, and monocytes (THP-1), muscle (RD) cells, and T lymphocytes (Jurkat, H9). The same pseudotype system was used to demonstrate that BASV G-mediated VSIV entry into adult grivet kidney (Vero), Asian tiger mosquito larva (C6/36, C7/10), aurochs (*Bos taurus*) kidney (MDBK), BHK-21, Brazilian free-tailed bat (*Tadarida brasiliensis*) lung (Tb1 Lu), brown rat (*Rattus norvegicus*) kidney (NRK) and brain (C6) cells, domestic pig (*Sus scrofa*) kidney (SK-RST) cells, and house mouse (*Mus musculus*) fibroblasts (3T3, MC57) (Steffen et al., 2013). These experiments suggested that BASV particles can enter a broad range of cells from diverse animal species, including *Homo sapiens*. This result stands in contrast to the knowledge available at the time on tropism of other tibroviruses.

Our systematic approach sheds further light on tibrovirion cell entry abilities. Together, our results indicate that all tibrovirus Gs, not only BASV G, can mediate tibrovirus particle entry into a wide variety of cells derived from animals of diverse species, including *Homo sapiens* (Figures 2, 3, and 5). The most surprising result was that EKV-1 and EKV-2, which had been discovered in human sera (Stremlau et al., 2015), appeared so much more efficient in entering human cells than BASV, the only other tibrovirus found in humans thus far (Grard et al., 2012) (Figure 2).

However, our results need to be regarded with caution. First, it is important to emphasize that our data only indicate which cell lines allow entry of particles of diverse tibroviruses, but we cannot state which of these tibrovirion-susceptible cells are also permissive to tibrovirus replication, particle formation, and particle egress. For instance, BASV, EKV-1, and/or EKV-2 could possibly enter numerous human cell types *in vivo* but then be restricted via antiviral pathways of the penetrated cells in replication and virion progeny formation. Indeed, N-detecting western blots using cells infected with four authentic tibroviruses (Figure 5) suggest that most cell lines are resistant to productive tibrovirus infection (absence of CPE and/or N expression) with the notable exception of TIBV. Vice versa, in Asian tiger mosquito (C6/36) cells, which lack a functional RNAi antiviral response (Brackney et al., 2010), all four authentic tibroviruses replicated efficiently. We measured N expression only at one timepoint due to the scale of the experiment, and we were unable to carefully quantify the titer and infectious-to-noninfectious particle ratios for the cell infection experiments. Hence, lack of N detection could be due to delayed or slow replication, whereas N detection could be partially due to detection of N stemming from incoming virions rather than from newly made transcripts. Second, the passaging histories of all isolated tibroviruses used in our study are complex, involving cells and animals of different species (Table 1). Unpassaged or low-passaged tibroviruses or tibrovirus sequences (or original samples) are not available anymore. Because the tibrovirus G sequences used for the creation of rVSIVs for this study were ultimately derived from these passaged viruses (Gubala et al., 2011; Lauck et al., 2015; Walker et al., 2015), we can only claim these those viruses enter animal cells efficiently, but we cannot extrapolate these results to wild-type tibroviruses. Interestingly, authentic TIBV entered (and likely infected) the majority of tested cell lines (Figure 5) largely in agreement with rVSIV-TIBV data (Figures 2 and 3). This result is somewhat puzzling as the closest relative of TIBV, BAV, which is classified in the same species as TIBV (Walker et al., 2015), did not cause CPE in most cell lines and BAV N could not be detected in them via western blotting, either. One explanation for this discrepancy is that the TIBV isolate we obtained evolved to be able to use mammalian and reptilian cells during repeated *in vitro* and *in vivo* passaging over the last decades, whereas BAV evolution took a different path because of a different passaging history (Table 1). In that regard it is worth mentioning that the experiment with BAV also produced individual puzzling results: for instance, three human cancer cell lines (melanoma M14, human colon cancer HCC-2998, renal cancer A498) developed CPE after BAV exposure but BAV N could not be detected; vice versa, four human cell lines (renal cancer ACHN, lung cancer A549, NCI-H23, and NCI-H460) did not develop CPE but BAV N could be detected. The latter result could indicate nondetrimental viral replication (or detection of incoming virion N), whereas the former result remains to be explained.

Together, the data obtained during our study indicate that tibrovirus tropism evaluations using tibrovirus surrogate systems (such as pseudotyped VSIVs or rVSIVs) should not be interpreted as indicative of the actual ability of a tibrovirus to infect, for instance, humans. However, these studies will be useful to further evaluate the general entry mechanism of tibroviruses, including, for instance, the identification of (the) tibrovirus cell surface receptor(s), which appear to be rather ubiquitously expressed. To evaluate the pathogenic potential of alleged human tibroviruses such as BASV, EKV-1, and EKV-2, replicating isolates will have to be obtained either by means of isolation from a natural host reservoir or vector or by means of reverse genetics.

## ACKNOWLEDGMENTS

We would like to thank Charles H. Calisher (Colorado State University, Fort Collins, CO, USA), Ania Gubala (CSIRO, Australia), and Toby St. George (Australia) for sharing knowledge and historic reports on tibroviruses, and Laura Bollinger (IRF-Frederick, Frederick, MD, USA) for critically editing the manuscript.

## AUTHOR CONTRIBUTIONS STATEMENT

YC, SY, ENP, ML, MRW, CLF, and SRR performed experiments. RKJ, SPJW, and KC developed and grew recombinant vesicular stomatitis Indiana viruses expressing tibrovirus G glycoproteins. RBT grew tibrovirus stocks. JW, DHO’C, GP, CYC, and JHK planned experiments and analyzed data. JHK wrote the manuscript. All authors reviewed and approved the manuscript.

## DISCLAIMER

The content of this publication does not necessarily reflect the views or policies of the US Department of the Army, the US Department of Defense, or the US Department of Health and Human Services or of the institutions and companies affiliated with the authors.

---

**Supplementary Figure 1.**
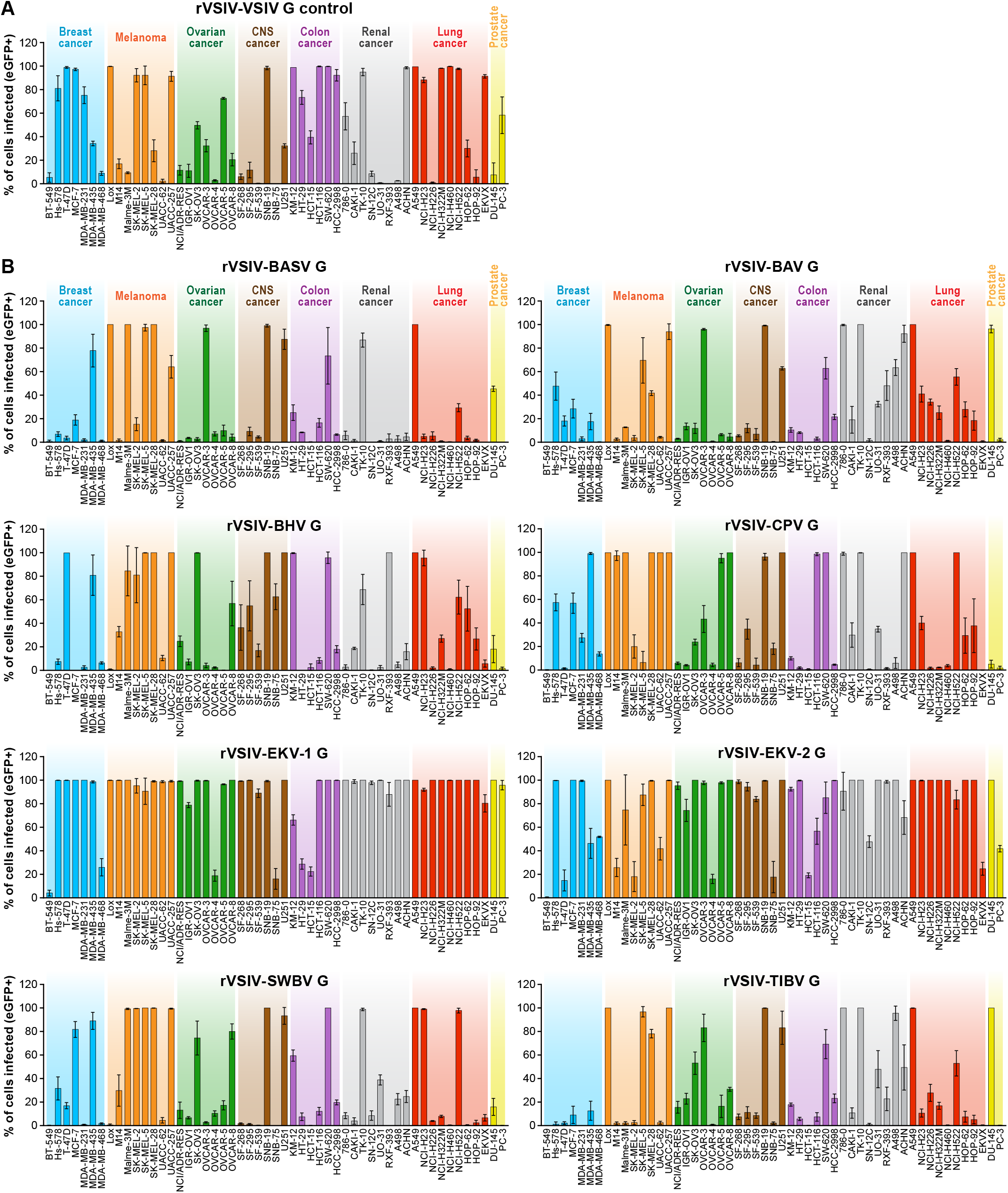
Tibrovirus glycoproteins mediate virion entry into a broad range of human cell types. Same experiment as in Figures 1B and 2 using rVSIVs expressing diverse tibrovirus glycoproteins (G) (MOI = 0.3). The percentage of eGFP-expressing NCI-60 human cell panel cell lines was measured by high-content imaging at 24 h post-exposure. All experiments were performed in triplicate; error bars show standard deviations. BHV, Beatrice Hill virus; BASV, Bas-Congo virus; BAV, Bivens Arm virus; CNS, central nervous system, CPV; Coastal Plains virus; EKV-1, Ekpoma virus 1; eGFP, enhanced green fluorescent protein; EKV-2, Ekpoma virus 2; MOI, multiplicity of infection; SWBV, Sweetwater Branch virus; TIBV, Tibrogargan virus; rVSIV, recombinant vesicular stomatitis Indiana virus. NCI-60 cell lines are listed by their abbreviations and grouped by organ/cancer type.

**Supplementary Figure 2.**
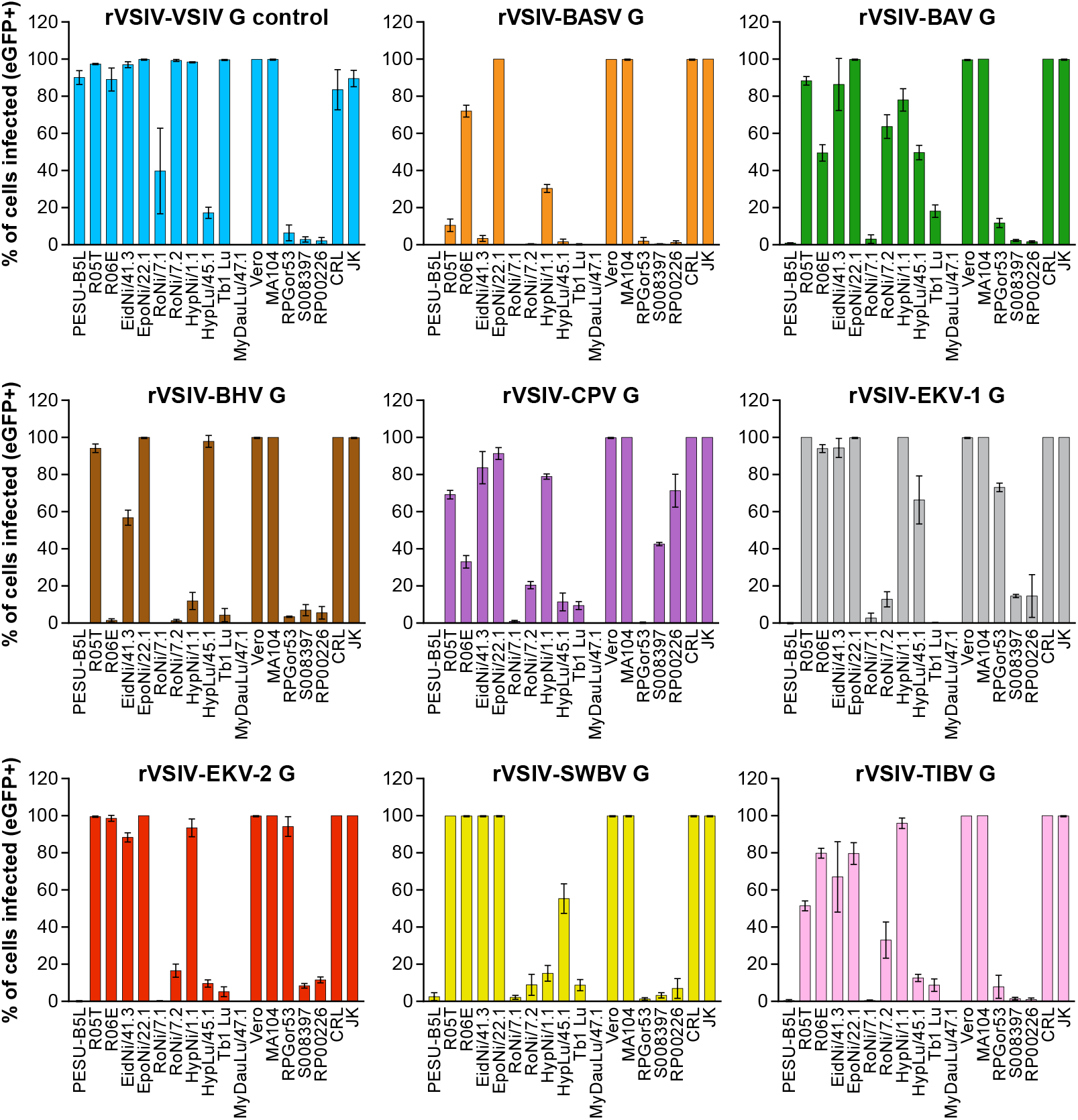
Tibrovirus glycoproteins mediate virion entry into a broad range of animal cell types. Same experiment as in Figure 3 using different cell types exposed to rVSIV-VSIV G control and rVSIVs expressing diverse tibrovirus glycoproteins (G) (MOI=0.3). **(A)** Bat (PESU-B5L, Ro5T, Ro6E, EidNi/41.3, EpoNi/22.1, RoNi/7.1, RoNi/7.2, HypNi/1.1, HypLu/45.1, Tb1 Lu, MyDauLu/47.1), nonhuman primate (Vero, MA104, RPGor53, S008397, RP00226), hispid cotton rat CRL, and boa constrictor JK cell lines. **(B)** Asian tiger mosquito C6/36 cells. The percentage of eGFP-expressing cell lines was measured by high-content imaging at 24 h post-exposure (bat, nonhuman primate, hispid cotton rat, and boa constrictor cell lines) or at 24 h, 48 h, 72 h, and 96 h post-exposure (Asian tiger mosquito cells). All experiments were performed in triplicate; error bars show standard deviations. BHV, Beatrice Hill virus; BASV, Bas-Congo virus; BAV, Bivens Arm virus; CPV, Coastal Plains virus; eGFP, enhanced green fluorescent protein; EKV-1, Ekpoma virus 1; EKV-2, Ekpoma virus 2; SWBV, Sweetwater Branch virus; TIBV, Tibrogargan virus; rVSIV, recombinant vesicular stomatitis Indiana virus.

## REFERENCES

Abraham, G., and Banerjee, A.K. (1976). Sequential transcription of the genes of vesicular stomatitis virus. Proc Natl Acad Sci U S A 73(5), 1504–1508.

Altstiel, L.D., and Landsberger, F.R. (1981). Lipid-protein interactions between the surface glycoprotein of vesicular stomatitis virus and the lipid bilayer. Virology 115(1), 1–9.

Babayan, S.A., Orton, R.J., and Streicker, D.G. (2018). Predicting reservoir hosts and arthropod vectors from evolutionary signatures in RNA virus genomes. Science 362(6414), 577–580. doi: 10.1126/science.aap9072.

Ball, L.A., and White, C.N. (1976). Order of transcription of genes of vesicular stomatitis virus. Proc Natl Acad Sci U S A 73(2), 442–446.

Bearzotti, M., Delmas, B., Lamoureux, A., Loustau, A.-M., Chilmonczyk, S., and Bremont, M. (1999). Fish rhabdovirus cell entry is mediated by fibronectin. J Virol 73(9), 7703–7709.

Biesold, S.E., Ritz, D., Gloza-Rausch, F., Wollny, R., Drexler, J.F., Corman, V.M., et al. (2011). Type I interferon reaction to viral infection in interferon-competent, immortalized cell lines from the African fruit bat *Eidolon helvum*. PLoS One 6(11), e28131. doi: 10.1371/journal.pone.0028131.

Blumenthal, R., Bali-Puri, A., Walter, A., Covell, D., and Eidelman, O. (1987). pH-dependent fusion of vesicular stomatitis virus with Vero cells. Measurement by dequenching of octadecyl rhodamine fluorescence. J Biol Chem 262(28), 13614–13619.

Bourhy, H., Cowley, J.A., Larrous, F., Holmes, E.C., and Walker, P.J. (2005). Phylogenetic relationships among rhabdoviruses inferred using the L polymerase gene. J Gen Virol 86(Pt 10), 2849–2858. doi: 10.1099/vir.0.81128-0.

Brackney, D.E., Scott, J.C., Sagawa, F., Woodward, J.E., Miller, N.A., Schilkey, F.D., et al. (2010). C6/36 *Aedes albopictus* cells have a dysfunctional antiviral RNA interference response. PLoS Negl Trop Dis 4(10), e856. doi: 10.1371/journal.pntd.0000856.

Buchko, G.W., Clifton, M.C., Wallace, E.G., Atkins, K.A., and Myler, P.J. (2017). Backbone chemical shift assignments and secondary structure analysis of the U1 protein from the Bas-Congo virus. Biomol NMR Assign 11(1), 51–56. doi: 10.1007/s12104-016-9719-2.

Calisher, C.H., Borde, G.E.N., Tuekam, T., and Gibbs, E.P.J. (1993). “Antibody to Bivens Arm virus in Trinidadian buffalo: a temporal and evolutionary link between Australia, Asia and the Caribbean?, ” in *Bovine ephemeral fever and related rhabdoviruses. 1st international symposium, bovine ephemeral fever and related rhabdoviruses*. ACIAR Proceedings. (Canberra, Australia: ACIAR), 80–83.

Cheng, C.Y., Shih, W.L., Huang, W.R., Chi, P.I., Wu, M.H., and Liu, H.J. (2012). Bovine ephemeral fever virus uses a clathrin-mediated and dynamin 2-dependent endocytosis pathway that requires Rab5 and Rab7 as well as microtubules. J Virol 86(24), 13653–13661. doi: 10.1128/jvi.01073-12.

Chiu, C., Fair, J., and Leroy, E.M. (2013). Bas-Congo virus: another deadly virus? Future Microbiol 8(2), 139–141. doi: 10.2217/fmb.12.145.

Cybinski, D.H., and Gard, G.P. (1986). Isolation of a new rhabdovirus in Australia related to Tibrogargan virus. Aust J Biol Sci 39(3), 225–232.

Cybinski, D.H., St. George, T.D., Standfast, H.A., and McGregor, A. (1980). Isolation of Tibrogargan virus, a new Australian rhabdovirus, from *Culicoides brevitaris*. Vet Microbiol 5, 301–308.

Finkelshtein, D., Werman, A., Novick, D., Barak, S., and Rubinstein, M. (2013). LDL receptor and its family members serve as the cellular receptors for vesicular stomatitis virus. Proc Natl Acad Sci U S A 110(18), 7306–7311. doi: 10.1073/pnas.1214441110.

Florkiewicz, R.Z., and Rose, J.K. (1984). A cell line expressing vesicular stomatitis virus glycoprotein fuses at low pH. Science 225(4663), 721–723.

Garbutt, M., Liebscher, R., Wahl-Jensen, V., Jones, S., Möller, P., Wagner, R., et al. (2004). Properties of replication-competent vesicular stomatitis virus vectors expressing glycoproteins of filoviruses and arenaviruses. J Virol 78(10), 5458–5465.

Gard, G.P., Shorthose, J.E., Weir, R.P., Walsh, S.J., and Melville, L.F. (1988). Arboviruses recovered from sentinel livestock in northern Australia. Vet Microbiol 18(2), 109–118.

Gaudin, Y., Ruigrok, R.W.H., Knossow, M., and Flamand, A. (1993). Low-pH conformational changes of rabies virus glycoprotein and their role in membrane fusion. J Virol 67(3), 1365–1372.

Gibbs, E.P., Calisher, C.H., Tesh, R.B., Lazuick, J.S., Bowen, R., and Greiner, E.C. (1989). Bivens Arm virus: a new rhabdovirus isolated from *Culicoides insignis* in Florida and related to Tibrogargan virus of Australia. Vet Microbiol 19(2), 141–150.

Grard, G., Fair, J.N., Lee, D., Slikas, E., Steffen, I., Muyembe, J.-J., et al. (2012). A novel rhabdovirus associated with acute hemorrhagic fever in Central Africa. PLoS Pathog 8(9), e1002924. doi: 10.1371/journal.ppat.1002924.

Gubala, A., Davis, S., Weir, R., Melville, L., Cowled, C., and Boyle, D. (2011). Tibrogargan and Coastal Plains rhabdoviruses: genomic characterization, evolution of novel genes and seroprevalence in Australian livestock. J Gen Virol 92(Pt 9), 2160–2170. doi: 10.1099/vir.0.026120-0.

Hastie, E., Cataldi, M., Marriott, I., and Grdzelishvili, V.Z. (2013). Understanding and altering cell tropism of vesicular stomatitis virus. Virus Res 176(1-2), 16–32. doi: 10.1016/j.virusres.2013.06.003.

Hoffmann, M., Müller, M.A., Drexler, J.F., Glende, J., Erdt, M., Gützkow, T., et al. (2013). Differential sensitivity of bat cells to infection by enveloped RNA viruses: coronaviruses, paramyxoviruses, filoviruses, and influenza viruses. PLoS One 8(8), e72942. doi: 10.1371/journal.pone.0072942.

Huang, B., Allcock, R., and Warrilow, D. (2016). Newly characterized arboviruses of northern Australia. Virol Rep 6, 11–17. doi: 10.1016/j.virep.2016.01.001.

Huynh, J., Li, S., Yount, B., Smith, A., Sturges, L., Olsen, J.C., et al. (2012). Evidence supporting a zoonotic origin of human coronavirus strain NL63. J Virol 86(23), 12816–12825. doi: 10.1128/jvi.00906-12.

Iverson, L.E., and Rose, J.K. (1981). Localized attenuation and discontinuous synthesis during vesicular stomatitis virus transcription. Cell 23(2), 477–484.

Jangra, R.K., Herbert, A.S., Li, R., Jae, L.T., Kleinfelter, L.M., Slough, M.M., et al. (2018). Protocadherin-1 is essential for cell entry by New World hantaviruses. Nature 563(7732), 559–563. doi: 10.1038/s41586-018-0702-1.

Jordan, I., Horn, D., Oehmke, S., Leendertz, F.H., and Sandig, V. (2009). Cell lines from the Egyptian fruit bat are permissive for modified vaccinia Ankara. Virus Res 145(1), 54–62. doi: 10.1016/j.virusres.2009.06.007.

Kleinfelter, L.M., Jangra, R.K., Jae, L.T., Herbert, A.S., Mittler, E., Stiles, K.M., et al. (2015). Haploid genetic screen reveals a profound and direct dependence on cholesterol for hantavirus membrane fusion. MBio 6(4), e00801. doi: 10.1128/mBio.00801-15.

Kühl, A., Hoffmann, M., Müller, M.A., Munster, V.J., Gnirß, K., Kiene, M., et al. (2011). Comparative analysis of Ebola virus glycoprotein interactions with human and bat cells. J Infect Dis 204 Suppl 3, S840–849. doi: 10.1093/infdis/jir306.

Lafon, M. (2005). Rabies virus receptors. J Neurovirol 11(1), 82–87. doi: 10.1080/13550280590900427.

Lauck, M., Yú, S., Caì, Y., Hensley, L.E., Chiu, C.Y., O’Connor, D.H., et al. (2015). Genome sequence of Bivens Arm virus, a tibrovirus belonging to the species *Tibrogargan virus (Mononegavirales: Rhabdoviridae)*. Genome Announc 3(2), e00089–00015. doi: 10.1128/genomeA.00089-15.

Liu, H., Liu, Y., Liu, S., Pang, D.-W., and Xiao, G. (2011). Clathrin-mediated endocytosis in living host cells visualized through quantum dot labeling of infectious hematopoietic necrosis virus. J Virol 85(13), 6252–6262. doi: 10.1128/jvi.00109-11.

Maes, P., Amarasinghe, G.K., Ayllón, M.A., Basler, C.F., Bavari, S., Blasdell, K.R., et al. (2019). Taxonomy of the order *Mononegavirales:* second update 2018. Arch Virol, In press.

Moreira, É.A., Locher, S., Kolesnikova, L., Bolte, H., Aydillo, T., García-Sastre, A., et al. (2016). Synthetically derived bat influenza A-like viruses reveal a cell type- but not species-specific tropism. Proc Natl Acad Sci U S A 113(45), 12797–12802. doi: 10.1073/pnas.1608821113.

Piccinotti, S., Kirchhausen, T., and Whelan, S.P.J. (2013). Uptake of rabies virus into epithelial cells by clathrin-mediated endocytosis depends upon actin. J Virol 87(21), 11637–11647. doi: 10.1128/jvi.01648-13.

Raaben, M., Jae, L.T., Herbert, A.S., Kuehne, A.I., Stubbs, S.H., Chou, Y.-Y., et al. (2017). NRP2 and CD63 are host factors for Lujo virus cell entry. Cell Host Microbe 22(5), 688–696.e685. doi: 10.1016/j.chom.2017.10.002.

Regan, A.D., and Whittaker, G.R. (2013). Entry of rhabdoviruses into animal cells. Adv Exp Med Biol 790, 167–177. doi: 10.1007/978-1-4614-7651-1_9.

Robinson, L.R., and Whelan, S.P.J. (2016). Infectious entry pathway mediated by the human endogenous retrovirus K envelope protein. J Virol 90(7), 3640–3649. doi: 10.1128/jvi.03136-15.

Shao, L., Zhao, J., and Zhang, H. (2016). Spring viraemia of carp virus enters grass carp ovary cells via clathrin-mediated endocytosis and macropinocytosis. J Gen Virol 97(11), 2824–2836. doi: 10.1099/jgv.0.000595.

St George, T.D. (1985). Studies on the pathogenesis of bovine ephemeral fever in sentinel cattle. I. Virology and serology. Vet Microbiol 10(6), 493–504.

Standfast, H.A., Dyce, A.L., St George, T.D., Muller, M.J., Doherty, R.L., Carley, J.G., et al. (1984). Isolation of arboviruses from insects collected at Beatrice Hill, Northern Territory of Australia, 1974-1976. Aust J Biol Sci 37(5-6), 351–366.

Steffen, I., Liss, N.M., Schneider, B.S., Fair, J.N., Chiu, C.Y., and Simmons, G. (2013). Characterization of the Bas-Congo virus glycoprotein and its function in pseudotyped viruses. J Virol 87(17), 9558–9568. doi: 10.1128/JVI.01183-13.

Stenglein, M.D., Sanders, C., Kistler, A.L., Ruby, J.G., Franco, J.Y., Reavill, D.R., et al. (2012). Identification, characterization, and *in vitro* culture of highly divergent arenaviruses from boa constrictors and annulated tree boas: candidate etiological agents for snake inclusion body disease. MBio 3(4), e00180–00112. doi: 10.1128/mBio.00180-12.

Stremlau, M.H., Andersen, K.G., Folarin, O.A., Grove, J.N., Odia, I., Ehiane, P.E., et al. (2015). Discovery of novel rhabdoviruses in the blood of healthy individuals from West Africa. PLoS Negl Trop Dis 9(3), e0003631. doi: 10.1371/journal.pntd.0003631.

Sun, X., Yau, V.K., Briggs, B.J., and Whittaker, G.R. (2005). Role of clathrin-mediated endocytosis during vesicular stomatitis virus entry into host cells. Virology 338(1), 53–60. doi: 10.1016/j.virol.2005.05.006.

Tani, H., Morikawa, S., and Matsuura, Y. (2011). Development and applications of VSV vectors based on cell tropism. Front Microbiol 2, 272. doi: 10.3389/fmicb.2011.00272.

The University of Texas Medical Branch (UTMB) (2018). World Reference Center for Emerging Viruses and Arboviruses. https://www.utmb.edu/wrceva/home.

Tuekam, T., Greiner, E.C., and Gibbs, E.P. (1991). Seroepidemiology of Bivens Arm virus infections of cattle in Florida, St. Croix and Puerto Rico. Vet Microbiol 28(2), 121–127.

Walker, P.J., Blasdell, K.R., Calisher, C.H., Dietzgen, R.G., Kondo, H., Kurath, G., et al. (2018). ICTV Virus Taxonomy Profile: *Rhabdoviridae*. J Gen Virol 99(4), 447–448. doi: 10.1099/jgv.0.001020.

Walker, P.J., Firth, C., Widen, S.G., Blasdell, K.R., Guzman, H., Wood, T.G., et al. (2015). Evolution of genome size and complexity in the *Rhabdoviridae*. PLoS Pathog 11(2), e1004664. doi: 10.1371/journal.ppat.1004664.

Weinstein, J.N. (2006). Spotlight on molecular profiling: “Integromic” analysis of the NCI-60 cancer cell lines. Mol Cancer Ther 5(11), 2601–2605. doi: 10.1158/1535-7163.mct-06-0640.

Weir, D.L., Laing, E.D., Smith, I.L., Wang, L.-F., and Broder, C.C. (2014). Host cell virus entry mediated by Australian bat lyssavirus G envelope glycoprotein occurs through a clathrin-mediated endocytic pathway that requires actin and Rab5. Virol J 11, 40. doi: 10.1186/1743-422x-11-40.

Wiley, M.R., Prieto, K., Blasdell, K.R., Caì, Y., Campos Lawson, C., Walker, P.J., et al. (2017). Beatrice Hill virus represents a novel species in the genus *Tibrovirus (Mononegavirales: Rhabdoviridae).* Genome Announc 5(4), e01485–01416. doi: 10.1128/genomeA.01485- 16.

Wong, A.C., Sandesara, R.G., Mulherkar, N., Whelan, S.P., and Chandran, K. (2010). A forward genetic strategy reveals destabilizing mutations in the Ebolavirus glycoprotein that alter its protease dependence during cell entry. J Virol 84(1), 163–175. doi: 10.1128/JVI.01832-09.

Wunner, W.H., Reagan, K.J., and Koprowski, H. (1984). Characterization of saturable binding sites for rabies virus. J Virol 50(3), 691–697.

